# Robust of AMBER force field parameters for glutathionylated cysteines

**DOI:** 10.1101/2023.09.18.558208

**Authors:** Zineb Elftmaoui, Emmanuelle Bignon

## Abstract

S-glutathionylation is an oxidative post-translational modification which is involved in the regulation of many cell signaling pathways. Increasing amounts of studies show that it is crucial in cell homeostasis and deregulated in several pathologies. However, the effect of S-glutathionylation on proteins structure and activity is poorly understood, and a drastic lack of structural information at the atomic scale remains. Studies based on the use of molecular dynamics simulations, which can provide important information about modification-induced modulation of proteins structure and function, are also sparse and there is no benchmarked force field parameters for this modified cysteine. In this contribution, we provide robust AMBER parameters for S-glutathionylation, that we tested extensively against experimental data through a total of 33 *μ*s molecular dynamics simulations. We show that our parameters set efficiently describe the global and local structural properties of S-glutathionylated proteins. These data provide the community with an important tool to stimulate investigations about the effect of S-glutathionylation on protein dynamics and function, in a common effort to unravel the structural mechanisms underlying its critical role in cellular processes.

## 1. Introduction

Among the many post-translation modifications (PTM) regulating cellular processes, protein S-glutathionylation (SSG) has a central role in redox signaling pathways yet remains largely understudied. This PTM consists in the addition of the tripeptide glutathione (GSH, a very abundant low-molecular weight thiol) to target cysteines, and acts as a key player in redox regulation of protein function in human, plants and bacteria [1] - see Figure 1.

**Figure 1.**
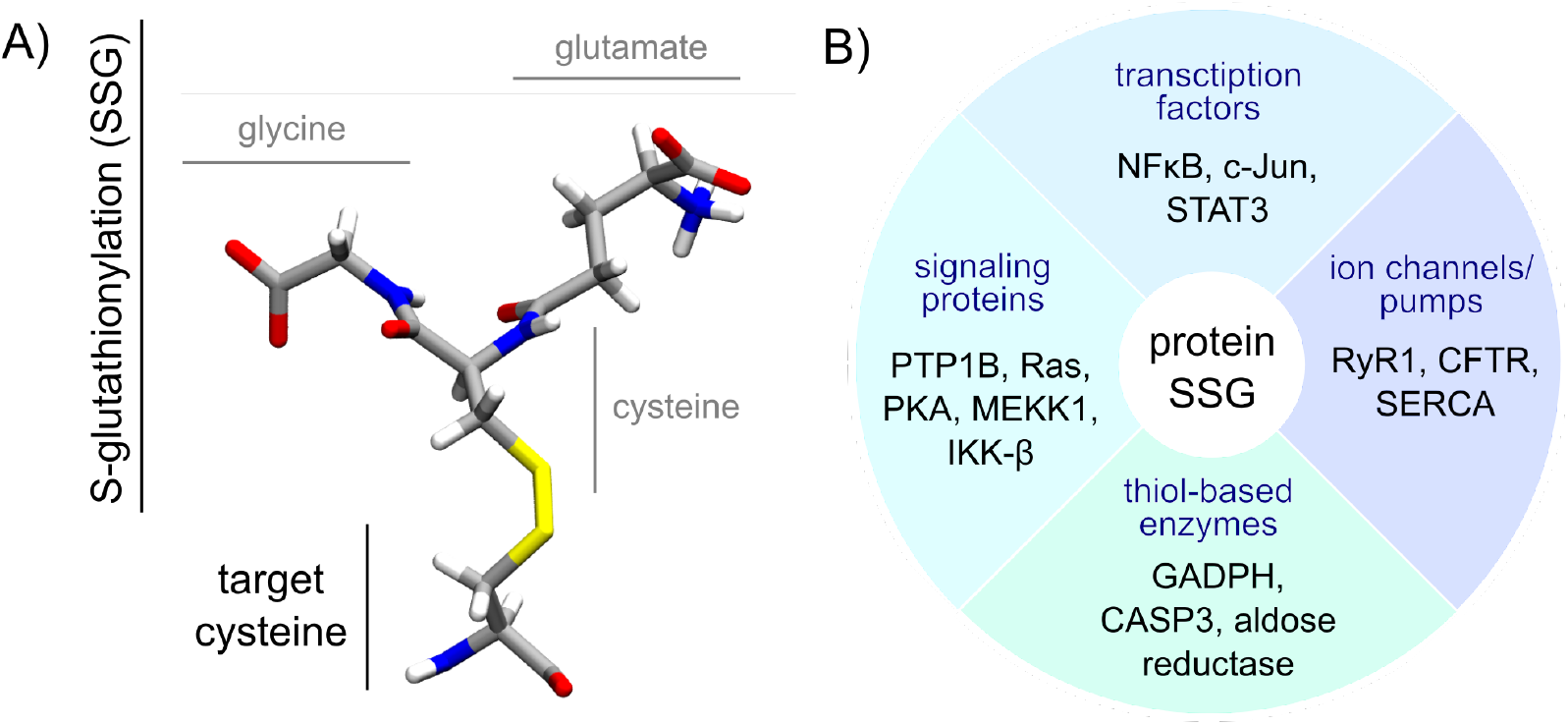
A) Structure of a glutathionyated cysteine. The glycine, cysteine and glutamate residues composing the glutathionyation moiety (SSG) are labeled in grey. The target cysteine backbone atoms are displayed as if embedded in a protein (without capping). B) Few examples of proteins regulated by S-glutathionylation, with respect to their function [8,16,17].

SSG participates to cellular homeostasis and has a crucial redox regulatory role in a large panel of cellular processes (e.g., apoptosis, proliferation, DNA compaction [2–5]). In response to an increase of Reactive Oxygen Species (ROS) amounts in the cell, SSG levels are upregulated to modulate the activity of a plethora of key-proteins, from signaling proteins to transcription factors and ion channels [6–8]). SSG is also thought to have a protective role against cysteine irreversible oxidation (e.g., sulfonylation) that could be generated under oxidative stress, and plays a crucial role in the regulation of proteins involved immune/inflammatory response to infection [9,10]. Elevated GSH and SSG levels have been observed in numerous diseases associated with oxidative stress and are most probably linked to anticancer drug resistance [11–14], further underlining the high biological relevance of this PTM in health and disease.

In cells, S-glutathionylation can be formed non-enzymatically [9]: by S-thiolation involving GSH and an oxidized cysteine, by S-thiolation of oxidized gluthatione with a cysteine, or by thiol/disulfide exchange between GSH and a cysteine thiolate group [13]. Although this modification was thought to occur in highly oxidative conditions, it is now well known that it happens in physiological conditions and is an important player in physiological redox signaling and cellular homeostasis. SSG levels are are also regulated by dedicated enzymes which act as writers and erasers of S-glutathionylated cysteines (CSG). Recent advances in experimental techniques dedicated to the study of this kind of reversible redox modifications brought out important information about SSG formation and removal [15], yet many aspects remain to be uncovered regarding this topic. Glutathione transferase enzymes (GST) have been shown to catalyze protein S-glutathionylation, and the glutaredoxin Grx1 is the main responsible for protein deglutathionylation in mammalian cells, with higher activity than the thioredoxin system [9].

Despite a growing interest of scientists in unraveling regulatory mechanisms of protein glutathionylation, the amount of all atom molecular dynamics studies present in the literature remains scarce compared to other mainstream PTM such as phosphorylation [18]. In most recent computational studies, Hyslop et al. reported the structural impact of SSG on the GADPH protein structure [19], Moffet et al. described the SSG-induced allosteric effects in a plant kinase [20], Zhou et al. investigated the effect of SSG on ATP/ADP binding to the adenine nucleotide translocator 2 protein [21], Zaffagnini et al. provided clues on SSG triggered collapse of a soluble tetrameric plant protein [22], and Hameed studied the inhibition of triose-phosphate isomerase by SSG [23]. However, CSG parameterization is rarely detailed, sometimes not even mentioned, and parameters sets are not provided. Yet, in order to ensure the reproductibility and relevance of the simulations of CSG-harboring systems, a thorough validation of CSG parameters against experimental data needs to be performed and made available to the community.

In order to palliate the absence of any extensively tested force field parameters for S-glutathionylated cysteines, we report here a set of AMBER parameters and their validation using eleven available experimental structures of S-glutathionylated proteins. Parameters were tested owing to microsecond-range MD simulations, performed starting either directly from the S-glutathionylated experimental structure, or from an Alphafold-predicted unmodified structure that was further S-glutathionylated *in silico*. Indeed, most of the future computational investigations of S-glutathionylated proteins will probably face the unavailability of any experimental starting structure for their simulations, hence the importance of testing the parameters set in similar conditions. We show that our AMBER-type parameters set succeeds at reproducing experimental global and local structural properties of S-glutathionylated proteins, both starting from experimental and *in silico* mutated structures. When starting from an *in silico* generated model of the modified protein, we advice to use multiple starting positions featuring different *χ* side chain angles for the S-glutathionylated cysteine. This work provides the community with a robust tool to foster new initiatives aiming at unveiling the role of protein S-glutathionylation in cells.

## 2. Materials and Methods

All MD simulations, QM calculations and structural analyses were performed with NAMD3 [24], Gaussian16 [25] and the Ambertools20 [26], respectively.

### 2.1. Generation of the CSG parameters

The inital geometry of an S-glutathionylated cysteine was taken from the CSG entry on the Protein Data Bank website, and capped with an acetyl (-OCH_3_) and a methylamino (-NHCH_3_) group on the N-term and C-term ends, respectively. This structure was optimized at the B3LYP/6-31+G* level, using a Polarizable Continuum Model to model an implicit solvent (water, *ϵ*=80), and setting the charge to -1 and the spin multiplicity to 1. Frequency calculation was performed in order to ensure the convergence of the minimization to a local potential energy well, characterized by the absence of negative vibrational frequency. The optimized geometry was extracted and subjected to a charge calculation using the Merz-Singh-Kollman scheme [27], at the HF/6-31+G* level.

The *antechamber* AMBER module was then used to extract the geometry, fit RESP charges, and assign atom types into a mol2 file. Atom names were kept as in the CSG entry of the PDB as much as possible. The charge of the capping atoms were set to 0, the backbone atom charges were set to match the cysteine charges from ff14SB [28], and the global charge difference was equally distributed on the side chain atoms (Δ of -0.0018 elementary charge per atom) to ensure a total residue charge of -1. The atom types were corrected to match these of ff14SB for the backbone, and let as guessed for the side chain after checking these were correctly assigned by *antechamber*. The *parmchk2* AMBER module was used to check the presence of missing parameters to describe the CSG residue, but none were identified. An AMBER library file for the CSG residue was finally created from the mol2 file and the ff14SB parameters using the *tleap* AMBER module, while removing the capping atoms and setting the amino acid connectivity. The parameter files are available on our Github: https://github.com/emmanuellebignon/CSG-parameters.

### 2.2. System setup

Each crystal structure was downloaded from the PDB and submitted to the propka3 software [29] for protonation state assignment. Crystallographic ions and waters were removed, and C- and N-terminal domains were reconstructed when needed - see details **in SI**. Besides, structures of the unmodified systems were generated using Alphafold2 through the Colabfold online facility [30,31], using the same primary sequence as the corresponding crystal structure. These initially unmodified structures were mutated *in silico* with *tleap* to retrieve the S-glutathionylated sites, in order to probe how robustly we could predict the experimental structures from a fully *in silico* model, which will mostly be the actual situation in further studies of S-glutathionylated proteins. Protonation states were assigned with respect to the propka3 results for the experimental structure.

Each system was soaked into a TIP3P[32] water box (truncated octahedron) of 12 Å buffer, neutralized with Na^+^and Cl^*−*^counter-ions [33]. Topology and coordinates parameters were generated using the *tleap*, using the ff14SB AMBER force field[28] and the CSG library designed in-house.

### 2.3. Molecular Dynamics simulations protocol

Each system was first minimized in 15,000 steps using the conjugate gradient method. The system was then thermalized to 300K and equilibrated in NPT along three runs of 600 ps with decreasing constraints on the solute. The time step was then changed from 2 fs to 4 fs by applying the Hydrogen Mass Repartioning scheme [34], in the 1*μ*s production runs. Three replicates were performed for each system, and the information and checkpoint files were outputted every 40 ps. The Particle Mesh Ewald approach [35] was applied for electrostatic calculations, with a 9 Å cutoff. The simulations were carried out in NPT, with a Langevin thermostat (1 ps^*−*1^ collision frequency) and a Langevin piston barostat.

MD trajectories were visualized and centered using the VMD software [36]. RMSD, radii of gyration, distances and dihedrals descriptors calculations, as well as the clustering analysis were performed using the *cpptraj* AMBER20 module. Statistical values were computed and plotted with the ggplot2 R package in Rstudio [37,38].

## 3. Results

### 3.1. Experimental structure curation

Reference structures of experimental S-glutathionylated proteins were curated from the Protein Data Bank by imposing a query on the GSH residue. Among the structures found, only these without any ligand, cofactor, other PTM, metal center, transmembrane domain, or internal missing region, were kept. Structure with a resolution higher than 2.0Å were evicted. This resulted in a selection of eleven proteins, mostly involved in redox processes such as thioredoxins, glutaredoxins and glutathione transferases besides the structure of the human lysosyme. All of these structures were obtained by X-Ray crystallography, and three of them are homodimers harboring two S-glutathionylation sites (TvGSTO2C, hGSTO2 and PtGSTF1). These structures are listed in Table 1 - see **SI** for more details.

**Table 1.** List of the curated experimental reference structures providing the protein name, the organism, the abbreviation used for the system in this study, the PDB ID of the structure, and the S-glutathionylation site.

| Protein name | Organism | Abbreviation | PDB ID | Resolution (Å) | Oxidation site |
| --- | --- | --- | --- | --- | --- |
| Thioredoxin 1 | *Saccharomyces cerevisiae* | ScTrx1 | 3F3R [39] | 1.80 | C30 |
| Thioredoxin-like2.1 | *Populus tremula x tremuloides* | PtTrxL2.1 | 5NYN [40] | 1.60 | C67 |
| Glutaredoxin S12 | *Populus tremula x tremuloides* | PtGrxS12 | 3FZ9 [41] | 1.70 | C29 |
| Glutaredoxin C5 | *Arabidopsis thaliana* | AtGrxC5 | 3RHB [42] | 1.20 | C29 |
| Glutaredoxin 2 | *Saccharomyces cerevisiae* | ScGrx2 | 3D5J [43] | 1.91 | C30 |
| Glutathione transferase λ1 | *Populus trichocarpa* | PtGSTL1 | 4PQH [44] | 1.40 | C36 |
| Glutathione transferase λ3 | *Populus trichocarpa* | PtGSTL3 | 4PQI [44] | 1.95 | C41 |
| Glutathione transferase ω2 | *Homo sapiens* | hGSTO2 | 3Q19<sup>1</sup> [45] | 1.90 | C29 |
| Glutathione transferase ω2C | *Trametes versicolor* | TvGSTO2C | 6SR8<sup>1</sup> [46] | 1.94 | C30 |
| Glutathione transferase F1 | *Populus tremula x tremuloides* | PtGSTF1 | 4RI7<sup>1</sup> [47] | 1.80 | C30 |
| Lysosyme | *Homo sapiens* | hLyso | 1HNL [48] | 1.40 | C95 |

<sup>1</sup> Homodimeric structures, harboring two symmetric oxidation sites.

For each of these systems, an unmodified model was generated *in silico* using the Alphafold-based Colabfold online tool. Noteworthy, the PDB file of the dimeric TvGSTO2C crystal structure (PDB ID 6SR8) displays a peculiar arrangement of the two monomers, different from the regular glutathione transferase quaternary structure - see Figure S1-A. Yet, as the model generated by Colabfold appears to exhibit a regular glutathione transferase fold, the system was simulated as such, and analyses were ran on separated monomers only. Besides, it is also important to underline that in the experimental structure of PtTrxL2.1, some atoms of CSG are missing (the terminal NH^+^ amino group of the side chain), presuming a high flexibility in this case - see Figure S1-B.

### 3.2. Global structural features of the systems

The global structural features of each system were calculated from the 3x1 *μ*s MD ensembles, in order to probe if our CSG parameter combined to ff14SB set would robustly describe the proteins 3D organization. For sake of clarity, we further refer to ‘crys’ and ‘AF’ simulations for the MD simulations from the crystal or from the Colabfold structures, respectively.

A first structural descriptor to scrutinize was the RMSD of each MD ensemble with respect to the reference experimental structure. We observed very good trends with average deviations systematically lower than 2.4 Å when considering all the atoms and lower than 1.6 Å for the backbone atoms - see Figure 2 and Figures S2 to S7.

**Figure 2.**
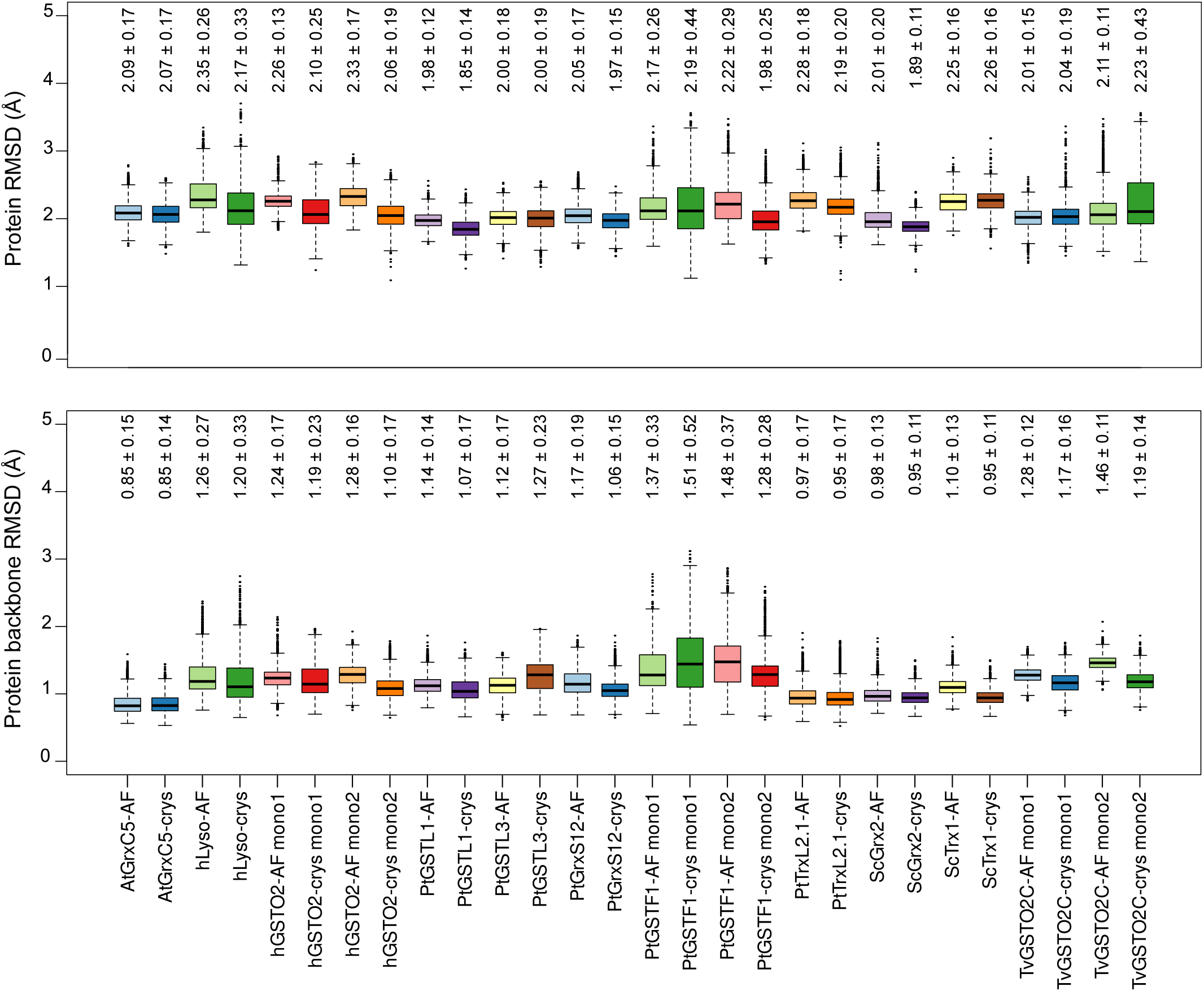
Boxplots of the RMSD values in Å of each system, considering all atoms (top) or the residues backbone atoms (bottom). RMSD values for dimers are shown by monomer (mono 1 or mono 2). The black lines inside each boxplot display the median values. Average and standard deviation values are written above each box.

Interestingly, the MD ensembles generated from the *in silico* models (AF simulations) exhibit average RMSD values within the range of the ones obtained for MD simulations starting from the experimental structure (crys simulations), both considering all the atoms or only the backbone atoms. Of note, the RMSD values of the dimers are of 2.36 *±* 0.22 Å and 2.22 *±* 0.28 Å for AF and crys PtGSTF1, and of 2.79*±* 0.33 Å and 2.20 *±* 0.23 Å for AF and crys hGSTO2 when considering all the atoms in the calculation - see Figures S2 and S5. The highest standard deviations are observed for the hLyso (up to 0.33 Å for all atoms), one PtGSTF1 monomer (up to 0.52 Å for backbone atoms) and one TvGSTO2C monomer (up to 0.43 Å for all atoms), which are mostly caused by fluctuations of some loops, as is evidenced from superimposing the representative structures of the major conformations identified by cluster analysis - see Figure 3. Of note, experimental resolutions were in a range of 1.2.-1.95 Å excluding hydrogen atoms, which underlines the robustness of our parameters combined to ff14SB in simulating the conformational behavior of these S-glutathionylated proteins.

**Figure 3.**
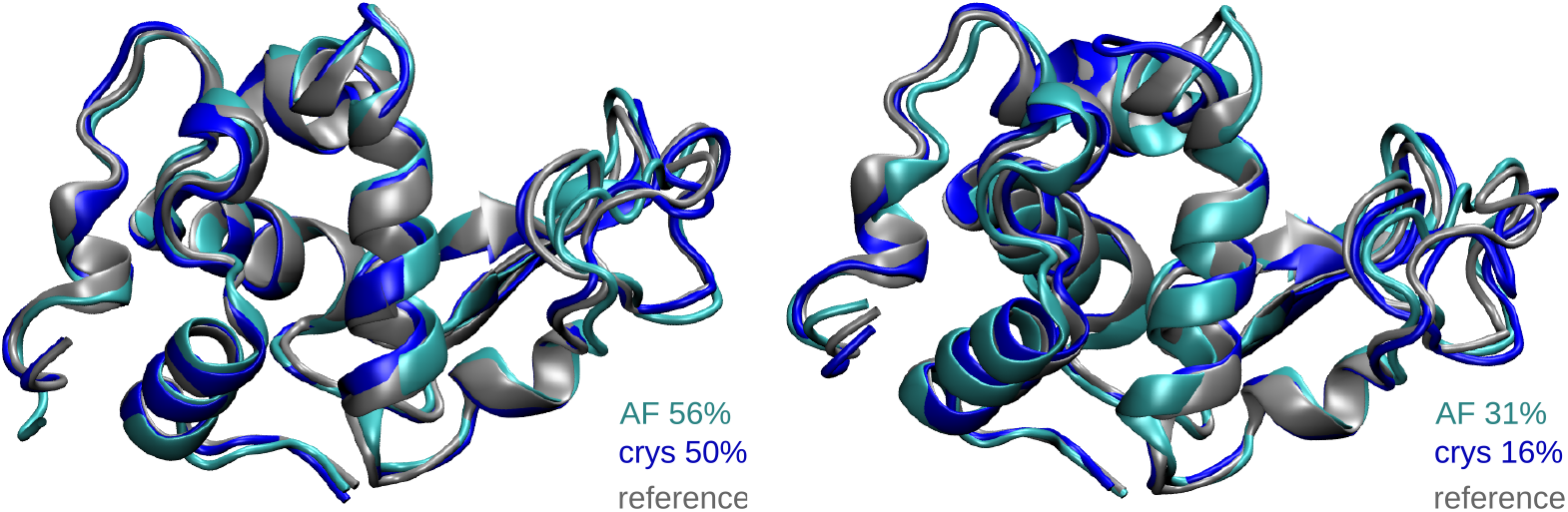
Representative structures of the two main conformations of hLyso as calculated from the cluster analysis of MD ensembles. Structures of the crystal reference and from the AF and crys simulations are displayed in gray, cyan and blue respectively. The percentage of occurrence of the clusters (primary on the left and secondary on the right) are also given.

**Figure 4.**
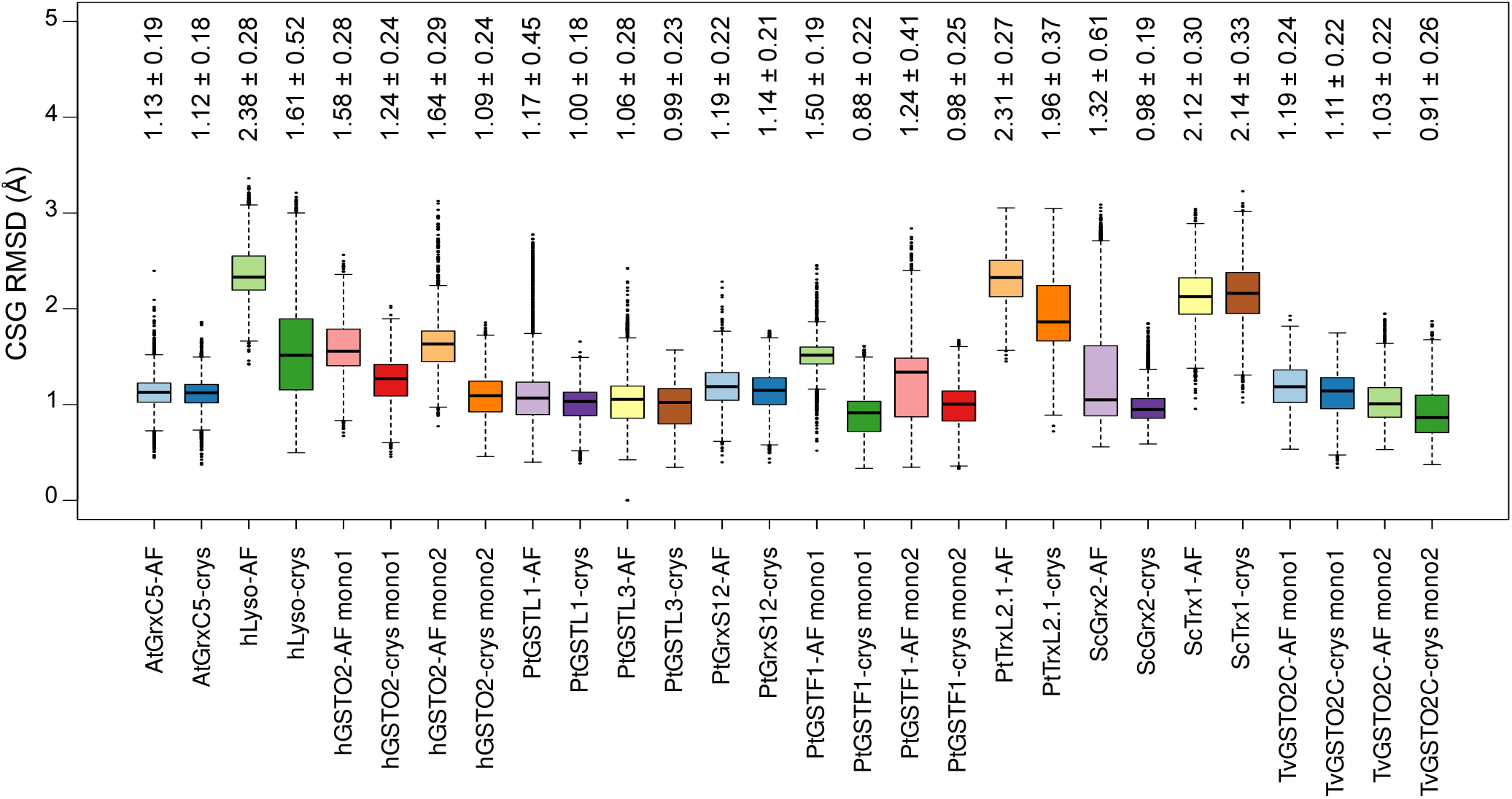
Boxplots of the S-glutathionylated cysteine (CSG) RMSD values in Å for each system. RMSD values for dimers are shown by monomer (mono 1 or mono 2). The black lines inside each boxplot display the median values. Average and standard deviation values are written above each box.

Another global descriptor of proteins is their radius of gyration, which provides information about their compactness. The average values calculated on the MD ensemble are in very good agreement with the reference values from the crystal structures, with a maximum deviation of only 0.4 Å observed for ScTrx1 AF simulations - see Table 2. Values obtained from the AF simulations are again within standard deviation of the ones calculated form the crys simulations, highlighting the robustness of our protocol. Noteworthy, values for TvGTO2C are not given considering the different quaternary structures of the AF and crys systems, as mentioned above.

**Table 2.** Average and standard deviation of the radius of gyration (Rg) for simulations from the crystal structure (crys) and from the Colabfold structure (AF). The reference value calculated from the experimental structure is also given. All values are in angström.

| System | Reference Rg | AF simulations Rg | Crys simulations Rg |
| --- | --- | --- | --- |
| AtGrxC5 | 13.2 | $13.3 \pm 0.1$ | $13.3 \pm 0.1$ |
| hLyso | 14.3 | $14.4 \pm 0.1$ | $14.4 \pm 0.1$ |
| hGSTO2 | 22.6 | $22.4 \pm 0.2$ | $22.7 \pm 0.2$ |
| PtGSTL1 | 16.9 | $17.2 \pm 0.1$ | $17.2 \pm 0.1$ |
| PtGSTL3 | 17.1 | $17.4 \pm 0.4$ | $17.4 \pm 0.1$ |
| PtGrxS12 | 13.2 | $13.4 \pm 0.1$ | $13.4 \pm 0.1$ |
| PtGSTF1 | 20.7 | $21.0 \pm 0.1$ | $21.0 \pm 0.1$ |
| PtTrxL2.1 | 13.5 | $13.7 \pm 0.1$ | $13.7 \pm 0.1$ |
| ScGrx2 | 13.0 | $13.4 \pm 0.1$ | $13.4 \pm 0.1$ |
| ScTrx1 | 12.4 | $12.8 \pm 0.1$ | $12.7 \pm 0.1$ |

### 3.3. CSG structural features

In order to have a thorough validation of our CSG parameters, an in-depth investigation of the modified cysteine sites structural features and their interaction networks was carried out.

The CSG RMSD trends are much more disparate than what is observed for the protein RMSD. While for most of the systems there is a very good agreement with the experimental reference, illustrated by average RMSD values lower than 1.5 Å, some cases step out with values up to 2.38 *±* 0.28 Å for hLyso AF simulations. PtTRXL2.1 and ScTrx1 systems also exhibit higher RMSD values than the others for both AF and crys simulations, which can be rationalized by a weak interaction network around the CSG residue which gives it great flexibility, as described below.

Noteworthy, the CSG RMSD time series of the AF simulations globally show that even when starting from an *in silico* generated system, the CSG parameterization allows to retrieve the CSG reference geometry, characterized by a drop of RMSD after some time of simulation - see Figures 5 and Figure S2-9. We can cite as stricking example, the cases of PtGSTL1 and ScGRX2, for which the RMSD values of CSG is up to 2.5-3.0 Å at the beginning of the AF simulations, and then drops below 1.0 Å. For AtGrxC5, PtGSTL3 and PtGrxS12 the RMSD of CSG at t=0 ns is around 2.0 Å, and drops rapidly around 1.0 Å. In the case of the PtGSTF1 dimer, exchanges between two conformations close to the reference are observed, characterized by RMSD values of around 1.0 Å for the first one and 1.5 Å for the second one. The second conformation is also sampled in the crys simulation, yet to a lesser extent - see Figure S5. This can be explained by small changes in the interaction network of the CSG site (especially sampled in the monomer 2), which is described below. Besides, the CSG RMSD values in hGSTO2 monomer 2 of the first AF simulation replicate (MD1) do not show any drop, contrarily to the other replicates (MD2 and MD3), explaining the slightly higher average value (1.58 *±* 0.28 Å) compared to the crys simulations (1.24 *±* 0.24 Å). This is also the case for the CSG site in hGSTO2 monomer 1, which RMSD drops from 2.0 to 1.5 Å after only 500 ns and 800 ns of simulation in replicates MD1 and MD2 respectively, hence exhibiting an average RMSD of 1.58 *±* 0.28 Å in AF simulations compared to 1.24 *±* 0.24 Å in crys simulations.

**Figure 5.**
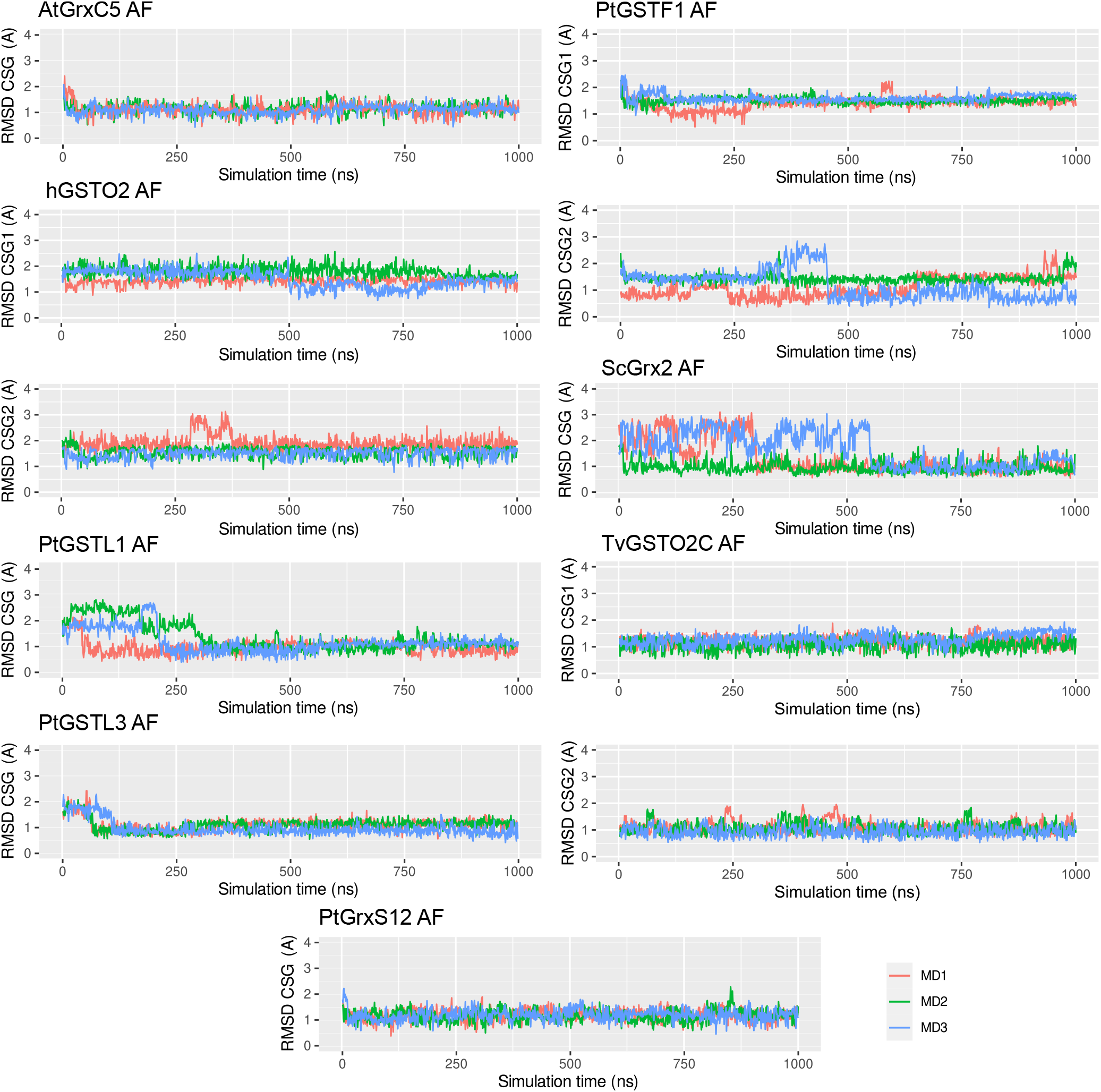
Times series of CSG RMSD values in Å of the three replicates for AF simulations of AtGrxC5, hGSTO2, PtGSTL1, PtGSTL3, PtGrx12, PtGSTF1, ScGrx2, and TvGSTO2C which show the better trends. Plots for the dimeric systems (hGSTO2, PtGSTF1 and TvGSTO2C) are paired in column, with CSG1 and CSG2 refering to monomer 1 and 2, respectively.

An important structural feature to check was the dihedral angle defining the new bond formed upon addition of the glutathione onto the cysteine residue, i.e. the dihedral angle around the S-S bond. Values from the AF and crys simulations were in excellent agreement with the reference structure, i.e. mainly +90^*°*^ - see Figure S10. PtGSTF1 is the only system where the CSG S-S angle is found at -90^*°*^ (only in monomer 1), which is accurately reproduced in both AF and crys simulations. We also note that values of this angle are rather centered around the average value, except for PtTrxL2.1 in which the distribution is broader for both AF and crys simulation yet still very much centered around the reference value of +90^*°*^. This might be due to the lack of strong hydrogen bonds involving the CSG side chain in this system (see details below), also resulting in higher CSG RMSD values in our simulations and in high B-factor values in the experimental structure (38.9 *±* 7.6 Å^2^ in average on the CSG atoms).

Concerning systems with the highest CSG RMSD deviations (i.e., hLyso, ScTrx1 and PtTrxL2.1), the superimposition of the major CSG conformations sampled suggest that the orientation of the CSG side chain in AF simulations is reversed with respect to the reference structure - i.e., the glutamate and the glycine sides are exchanged, see Figure 6-A. However, this does not explain why a high average of CSG RMSD is also observed in the crys simulations, which might be caused by a weak interaction network around the CSG, as described below. The RMSD time series show a conformational drift in PtTrxL2.1 crys simulations with an increase of RMSD from 1.7 Å to 2.3 Å in replicates MD1 and MD3. A similar behavior is observed in hLyso crys simulations with exchanges between several states in all replicates. Finally, CSG RMSD values in ScTrx1 AF and crys simulations are found to fluctuate a lot (2.12 *±* 0.30 Å and 2.14 *±* 0.33 Å respectively), without showing any stable conformation. An in-depth look at the interaction network involving the CSG residues helped to rationalize these observations.

**Figure 6.**
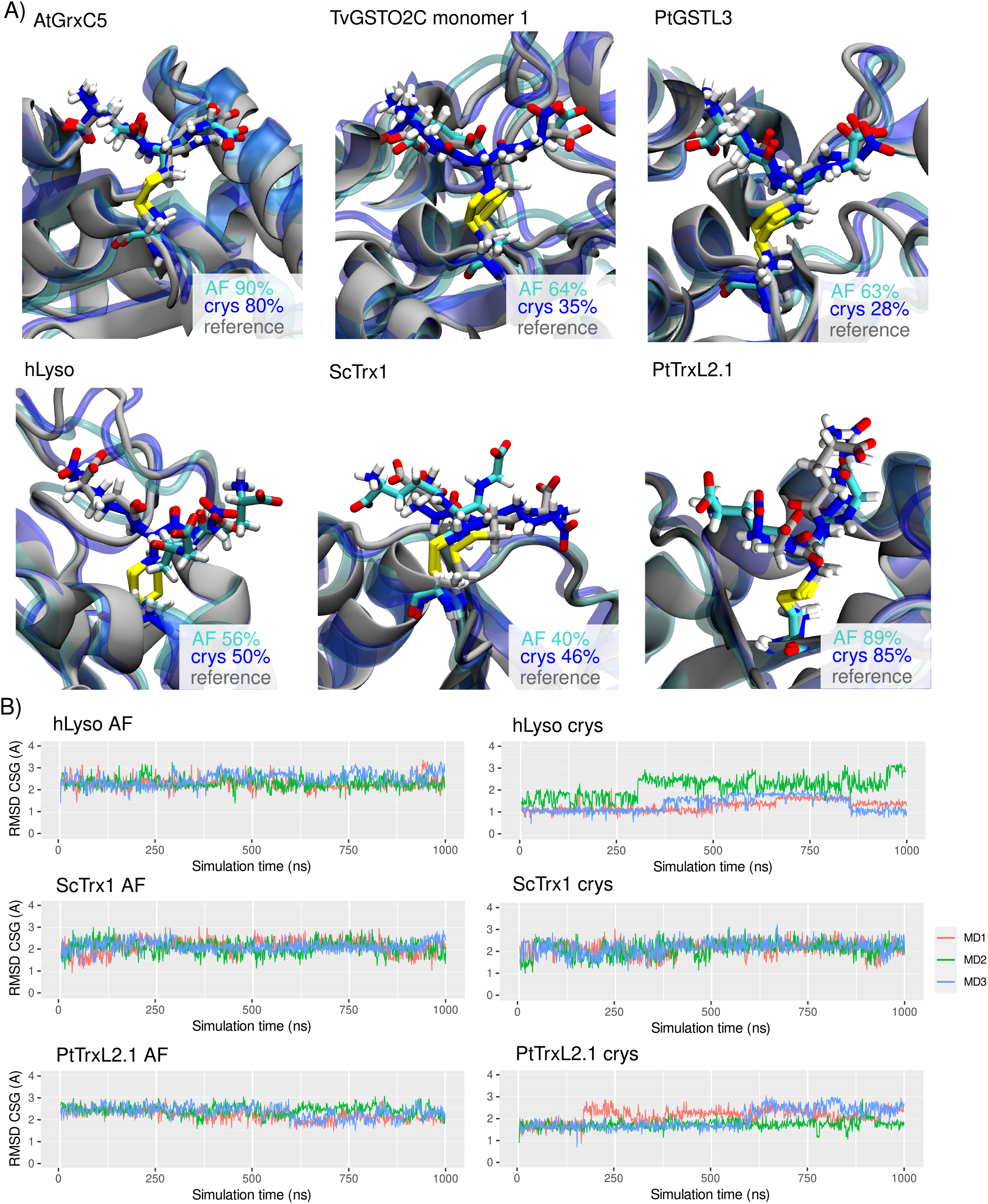
A) Representative structures of the major conformations of CSG in AtGrxC5, TvGSTO2C monomer 1, PtGSTL3, hLyso, ScTrx1, and PtTrxL2.1 as calculated from the cluster analysis of MD ensembles. Structures of the crystal reference and from the AF and crys simulations are displayed in gray, cyan and blue respectively. The percentage of occurrence of the clusters are also given. B) Times series of CSG RMSD values in Å of the three replicates for AF (left) and crys (right) simulations of hLyso, ScTrx1, and PtTrxL2.1.

### 3.4. CSG interaction network

Native interactions with the CSG side chain were listed for each system (see Table S2) and monitored along the AF and crys simulations. The CSG moieties involved in hydrogen bonds with the surroundings are the atoms of the amide moieties that results from dehydration steps of GSH formation (O2, OE1, H4 and H2), the carboxylate of the glycine end (C3) and the carboxylate and amino groups of the glutamate end (C1 and N3) - see Figure 7-A. Interestingly, conserved interaction patterns can by pinpointed in glutaredoxins which involve a valine snd a serine or threonine backbone atoms (V74 and T88 in both AtGrxC5 and PtGrxS12, and V75 and S89 for ScGrx2). Similar patterns are also found in the glutathione transferase proteins, with interactions between CSG.O2 and a valine backbone atoms (PtGSTL1 V79, PtGSTL3 V84, hGSTF1 V56, TvGSTO2C V56), between CSG.C1 and a serine (PtGSTL1 S92, PtGSTL3 S97, hGSTF1 S69, TvGSTO2C S81), and between CSG.N1 and a glutamate (PtGSTL1 E91, PtGSTL3 VE96, hGSTF1 E68, TvGSTO2C E80). hGSTO2 exhibits homologous interactions for CSG.O2 and CSG.C1 with I72 and S96, respectively. Consistently with the CSG RMSD trends, simulations results are in excellent agreement with the experimental reference. Most of the native interactions are observed in our simulations, and AF models generally succeed in retrieving the hydrogen bond (HB) network identified in the experimental structures - see Figures S11 to S15.

**Figure 7.**
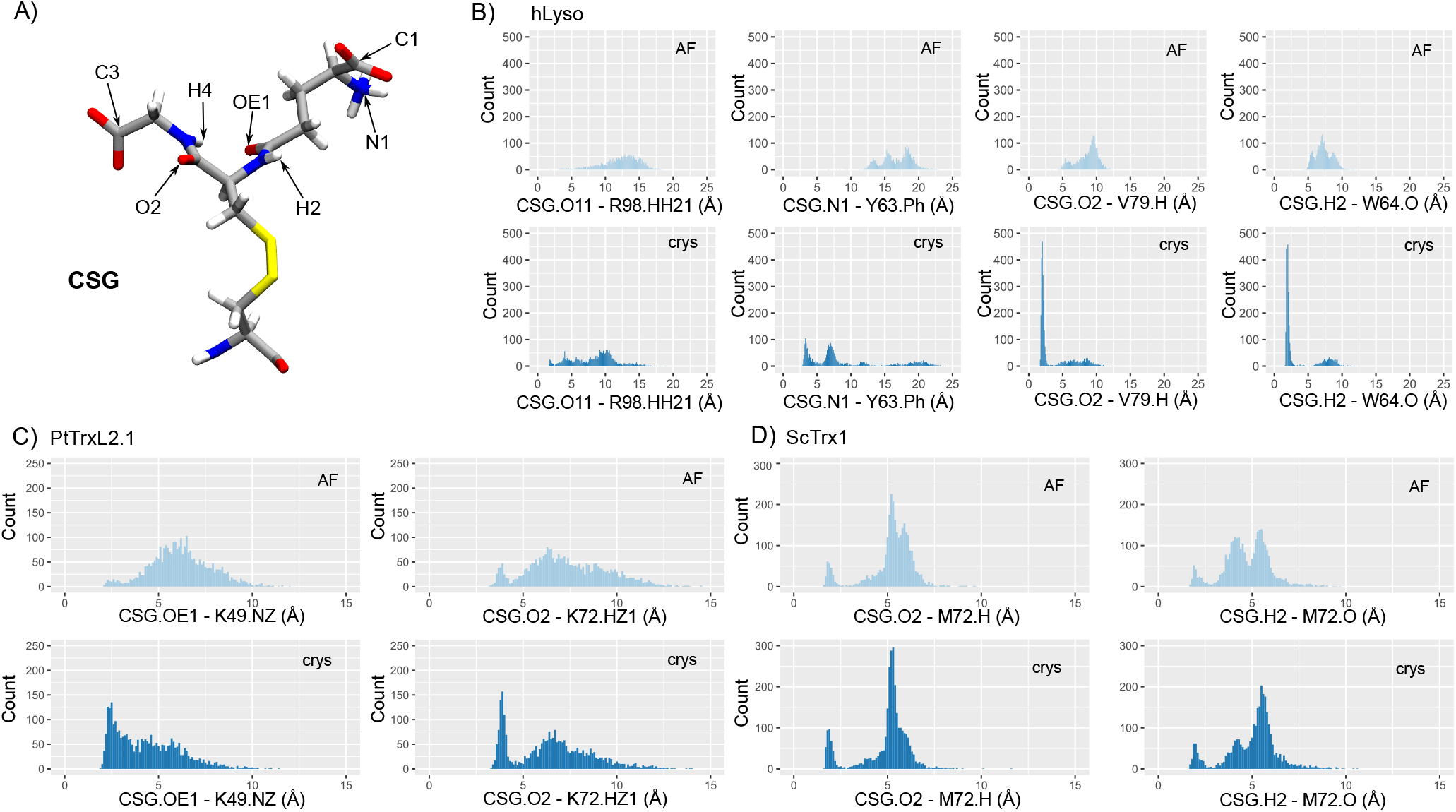
A) CSG structure, highlighting the side chain atoms used to monitor hydrogen bonds. B), C) and D) Distribution of distances corresponding to the native interactions identified in the reference structures of hLyso, PtTrxL2.1 and ScTrx1, respectively. Both distributions for AF (cyan) and crys (blue) simulations are shown. Cation-*π* interactions involving Y63 in hLyso are monitored as distances between CSG.N1 and the center of mass of the aromatic ring heavy atoms of Y63 (Y63.Ph).

All native hydrogen bonds are very efficiently reproduced in AtGrxC5, PtGSTL1, PtGSTL3, PtGrxS12, ScGrx2 and TvGTO2C, for both crys and AF simulations. Distance values in AF simulations of hGSTO2 and PtGSTF1 show broad distributions of values, probably because of the long simulation time necessary to retrieve the native CSG orientation - see Figures S2 and S5. As described previously in the CSG RMSD plot analysis, in some MD replicates the CSG geometry evolves to match the reference only in the late moments of the simulation. This is the case for hGSTO2 monomer 1, and in one of the replicates (MD1) the CSG conformation in monomer 2 does not match exactly the native orientation. Indeed, in this replicate it fails at retrieving the experimental HB network, especially the interaction between CSG.O2 and I72.H in both monomers. In monomer 1 only, the CSG.C1 - S96.HG hydrogen bond is not observed, yet CSG.C1 still strongly interacts with S96.H, suggesting that this is the major interaction for this site (also observed in the other glutathione transferase systems). The same trends are found for PtGSTF1, for which the CSG can adopt the same orientation as in the experimental structure but the HB network can deviate. Interactions for monomer 2 tend to exhibit distributions with peaks matching the reference, but this is less pronounced for monomer 1. Yet, the systematic presence of matching peaks suggests that these conformations are actually sampled, and the system might need more simulation time to stabilize itself. While AF simulations might require more extended sampling, results for crys simulations show in any case very robust trends in agreement with the reference.

The hLyso is a peculiar case, because it shows good results for the crys simulations while the AF simulations fail to correctly reproduce the native HB network - see Figure 7-B. In fact, the starting structure generated *in silico* features a CSG side chain with an orientation opposite to what is found in the reference structure. During the simulation, CSG is stuck in this conformation, and can not rotate to retrieve the native interactions that are found stable in the crys simulations, especially for CSG.O2 and H2 backbone atoms. We suggest that starting from multiple structures with opposite orientations of CSG side chain or resorting to enhanced sampling methods would be good solutions to overcome this problem.

Noteworthy, the native interactions in PtTrxL2.1 and ScTrx1 are found to be rather unstable in our simulations, with conformational fluctuations of the CSG side chain as suggested by the CSG RMSD values - see Figure 7-C and D. As a matter of fact, the only native interactions found for these two systems are a couple of HB involving the OE1, O2 or H2 atoms at the center of CSG side chain. No strong HB involving the CSG charged termini can be observed, probably because CSG is highly solvent-exposed in these two systems.

## 4. Discussion

Force field parameters for S-glutathionylated cysteines were generated and extensively tested against data from eleven curated experimental proteins bearing S-glutathionylation. Molecular Dynamics simulations of two types of starting systems were conducted: i) directly from the crystal structure or ii) from a 3D geometry predicted by the Alphafold-based Colabfold facility and S-glutathionylated *in silico* using the leap module of AMBER20. As only few experimental structures of S-glutathionylated proteins are available on the PDB, it was important to test our parameters on a fully *in silico* starting structure which might reflect the most probable situation encountered in future studies involving this modification.

The CSG parameters set was generated by calculating the RESP charges and assigning atom types from ff14SB. All the needed parameters to describe the CSG were already available in the ff14SB force field, and no specific parameterization was required. The CSG atom names were assigned to match the ones of the GSH entry on the Protein Data Bank, in order to facilitate their use for other systems. The mol2 and AMBER library files are available online (https://github.com/emmanuellebignon/CSG-parameters).

An extensive validation of the CSG parameters was then conducted. Eleven reference structures were identified in the PDB (see Table 1, which mostly featured redox-related proteins such as glutaredoxins and glutathione transferases. Three replicates of 1 *μ*s were carried out from the two types of starting systems, for each of the curated protein. The simulations starting from the crystal structure (crys simulations) generated conformations in excellent agreement with the reference data, both in terms of global (protein) and local (CSG site) features. The protein RMSD was systematically found below 2.3 Å when considering all the atoms (including hydrogens), and below 1.5 Å when taking into account only the backbone atoms - see Figure 2. The overall performances of our parameters were very good considering the resolution of the chosen reference structures, ranging from 1.20 to 1.95 Å. Importantly, conformations of the dimeric systems bearing two CSG sites were also very well reproduced in the crys simulations. The CSG features were also efficiently reproduced in the crys simulations, with RMSD values below 1.3 Å for most of the systems. Higher deviations were observed for the PtTrxL2.1 (1.96 *±* 0.37 Å) and ScTrx1 (2.14 *±* 0.33 Å) systems, mostly due to the weak interaction network around the CSG residue, resulting in a high flexibility of its side chain. The hLyso also exhibited slightly higher RMSD values for CSG (1.61 *±* 0.52 Å), which can be rationalized by the conformational exchanges in the simulations due to the movements of the disordered loop facing the CSG residue.

Simulations from the *in silico* starting structures provided more contrasted results. If the results show very good trends for most of the systems, with global and local features rapidly converging towards the reference values, some cases showed that such scenario must be taken with care. The most striking example illustrating this is the hLyso case, for which AF simulations failed to reproduced the CSG local features of the experimental structure - see Figures 6 and 7. This was due to the fact that the initial orientation of the CSG side chain is reversed with respect to the reference one. While in other systems the CSG reference geometry and interaction network are retrieved after some sampling time (see Figures S2 to S9), for hLyso the side chain is stuck in the initial wrong conformation in all replicates. We suggest that when starting from an *in silico* structure, multiple replicates should be launched from geometries featuring different CSG side chain orientations, or eventually to use enhanced sampling methods in order to boost the sampling of different conformations for the CSG residue. Given the excellent results obtained for the crys simulations, one should manage to converge to similar results in AF simulations by overcoming this starting orientation/sampling problem.

## 5. Conclusions

In conclusion, the combination of our CSG parameters with the ff14SB AMBER force field succeeded to reproduce the experimental global and local features of the experimental S-glutathionylated references. Simulations from the crystal structures provided conformations in excellent agreement with the reference data. A special care must be given when starting from *in silico* generated structures, as the CSG side chain can be stuck in a wrong initial conformation. We advise to multiply the MD replicates with different CSG orientations, in order to efficiently probe its conformational behavior. The use of enhanced sampling methods should also allow to overcome this issue. This AMBER parameters set for CSG is a robust tool for future computational investigations of S-glutathionylated systems, which will be of great importance towards the understanding of S-glutathionylation mechanisms of action as a redox post-translational regulator, and its link to disease onset and progression.

## Supporting information

Sup_Mat

## Supplementary Materials

The following supporting information can be downloaded at: https://www.mdpi.com/article/10.3390/ijms1010000/s1. Details of the references structures; Table S1: Residue ranges excluding N- and C-termini as used in the RMSD calculations and clustering analysis for each system; Table S2: Native interactions involving CSG side chain atoms as identified from the reference experimental structures; Figure S1: Quaternary structure of TvGSTO2C and CSG structure in PtTrxL2.1; Figures S2 to S7: RMSD time series; Figures S8 and S9: Superimposition of CSG main conformations with the reference structure; Figure S10: boxplot of the S-S dihedral angle; Figures S11 to S15: Distribution of the distances corresponding to native hydrogen bonds.

## Author Contributions

Conceptualization, E.B.; methodology, Z.E and E.B.; data curation, Z.E. and E.B.; writing—original draft preparation, Z.E and E.B.; writing—review and editing, Z.E and E.B.

## Funding

This research received no external funding.

## Data Availability Statement

All MD input files and post-processing scripts are available on Github. https://github.com/emmanuellebignon/CSG-parameters

## Acknowledgments

The authors thank GENCI and Explor computing centers for computational resources.

## Conflicts of Interest

The authors declare no conflict of interest.

## Disclaimer/Publisher’s Note

The statements, opinions and data contained in all publications are solely those of the individual author(s) and contributor(s) and not of MDPI and/or the editor(s). MDPI and/or the editor(s) disclaim responsibility for any injury to people or property resulting from any ideas, methods, instructions or products referred to in the content.

## Notes

https://github.com/emmanuellebignon/CSG-parameters

### Competing Interest Statement

The authors have declared no competing interest.

