## Supplementary material for "Robust of AMBER force field parameters for glutathionylated cysteines": Sup_Mat

### Reference structures details

##### Thioredoxins

Thioredoxins are ubiquitous enzymes that play a key role in cellular homeostasis for their antioxidant function. The thioredoxin superfamily gathers oxidoreductase enzymes with a common fold consisting in a four-strand $\beta$-sheets flanked by five $\alpha$-helices. One thioredoxin and one thioredoxin-like proteins belong to the set of reference structures used in this work. The S-glutathionylated structure of *Saccharomyces cerevisiae* thioredoxin Trx1 (ScTrx1) was obtained by Bao et al. in 2009, at a 1.80Å resolution (PDB ID 3F3R).[1] The modification site is on the catalytic C30 (C36 in the structure with the histone tag), and a C33S mutation was imposed to avoid internal di-sulfide bridge formation instead of S-glutathionylation. The Thioredoxin-like2.1 from *Populus tremula x Populus tremuloides* (PtTrxL2.1) is an atypical thioredoxin isoform that uses glutathione for regeneration. Its S-glutathionylated structure was published at a 1.20Å resolution by Chimani et al. in 2018 (PDB ID 5NYN),[2] with the modification site on the catalytic C67 residue. Noteworthy, the side chain of CSG was not complete for this structure (see Figure S1-B).

##### Glutaredoxins

Glutaredoxins are small oxidoreductase enzymes involved in the reduction of mixed disulfides in S-glutathionylated proteins. As they belong to the thioredoxin superfamily, they share the same fold. Three of the reference structures curated are glutaredoxins.

In 2009, Couturier et al. published the structure of the S-glutathionylated poplar glutaredoxin GrxS12 at a 1.7Å resolution (PtGrxS12, PDB ID 3FZ9).[3] The active site cysteine (C29) was the oxidation site. The same group resolved in 2011 the crystal structure of the monomeric apo-form of glutaredoxin C5 from *Aradobidopsis thaliana* (AtGrxC5) at a 1.20Å resolution (PDB ID 3RHB),[4] that features a S-glutathionylated cysteine at location 29 (active site). The S-glutathionylated structure of glutaredoxin Grx2 from the yeast *Sac-*

*charomyces cerevisiae* was released by Disola et al. in 2009[5] (ScGrx2, PDB ID 3D5J), with a 1.91Å resolution. The S-glutathionylation site is located on the catalytic C27 (PDB sequence C30), and the structure was obtained by mutating the nearby C30 (PDB sequence C33) to serine to avoid internal S-S bond formation.

##### Glutathione transferases

Glutathione transferases (GST) catalyze the conjugation of glutathione to proteins. Diverse types of GST exist in both prokaryotes and eukaryotes. Some of them are functional as monomers and others as dimers. Five of the curated reference structures are GST, two as monomers and three as dimers.

The two monomeric GST are Lambda glutathione transferases from Populus trichocarpa, published by Lallement et al. in 2014.[6] GST L1 (PtGSTL1, PDB ID 4PQH) and GST L3 (PtGSTL3, PDB ID 4QI) were resolve at a 1.4 and 1.95 Å resolution, respectively. Their fold is similar to these of Omega GST, yet the latter are functional as dimers while PtGSTL1/3 are strictly monomeric. They are S-glutathionylated on a conserved cysteine, C36 in PtGSTL1 and C41 in PtGSTL3.

Three dimeric GST have been selected from the PDB as reference structures. The first one is the human GST Omega 2 (hGSTO2, PDB ID 3Q19) published by Zhou et al. in 2012[7] at a 1.9 Å resolution, harboring S-glutathionylated cysteines symmetrically on C32 of both monomers. The structure of a Phi class GST from poplar (PtGSTF1, PDB ID 4RI7), released by Pegeot et al. in 2015[8] with a 1.80 Å resolution was also selected as reference. This structure also exhibit two symmetrical S-glutathionylation sites, on C13. The third dimeric GST, the GST Omega 2C from Trametes versicolor (TvGSTO2C, pDB ID 6SR8), also harbors two S-glutathionylated C13. It was published by Perrot et al.[9] in 2019, at a resolution of 1.94 Å. Strangely, the quaternary structure of TvGSTO2C as downloaded from the PDB does not exhibit a GST fold (yet the 3D view on the PDB) - see Figure S1-A. This structure was anyways kept as such for crys simulations, while AF simulations starting structure exhibited a standard GST geometry. Consequently, structural analyses were performed solely on the separated monomers.

**Human lysosyme**

The only experimental reference without any oxidoreductase function was the S-glutathionylated human lysosyme structure, reported by Inaka et al. in 1995 at a 1.80Å resolution (hLyso, PDB ID 1HNL).[10] A C77A mutation was introduced in order to avoid disulfide bridge with C95, the S-glutathionylated residue. The other cysteines of the protein are all involved in disulfide bridges (C6-C128, C30-C116, C65-C81).

### Supplementary figures and tables

Table S1 – Residue ranges excluding N- and C-termini as used in the RMSD calculations and clustering analysis for each system.

| System | Full range | No-termini range |
|---|---|---|
| ScTrx1 | 1-109 | 8-109 |
| PtTrxL2.1 | 1-122 | 11-122 |
| PtGrxS12 | 1-112 | 5-106 |
| AtGrxC5 | 1-113 | 6-105 |
| ScGrx2 | 1-112 | 6-109 |
| PtGSTL1 | 1-236 | 28-219 |
| PtGSTL3 | 1-241 | 33-223 |
| hGSTO2 | 1-478 | 24-231, 263-470 |
| TvGSTO2C | 1-490 | 5-236, 251-481 |
| PtGSTF1 | 1-430 | 5-214, 220-429 |
| hLyso | 1-130 | 5-128 |

Table S2 – Native interactions involving CSG side chain atoms as identified from the reference experimental structures

| **System** | **with CSG.O2** | **with CSG.H2** | **with CSG.OE1** |
|---|---|---|---|
| ScTrx1 | M78 | M78 | / |
| PtTrxL2.1 | K72 | / | K49 |
| PtGrxS12 | V73 | V73 | / |
| AtGrxC5 | V74 | V74 | / |
| ScGrx2 | V78 | V78 | / |
| PtGSTL1 | V79 | V79 | / |
| PtGSTL3 | V84 | V84 | / |
| hGSTO2 | I72 | I72 | / |
| TvGSTO2C | V56 | V56 | / |
| PtGSTF1 | V56 | V56 | / |
| hLyso | V79 | V79 | / |
| **System** | **with CSG.C1** | **with CSG.N1** | **with CSG.C3** |
| ScTrx1 | / | / | / |
| PtTrxL2.1 | / | / | / |
| PtGrxS12 | C86, T87 | / | / |
| AtGrxC5 | T88 | / | / |
| ScGrx2 | S92 | / | K27, E66 |
| PtGSTL1 | S92 | E91D | R65 |
| PtGSTL3 | S97 | E96 | R70, K83 |
| hGSTO2 | S96 | / | K59 |
| TvGSTO2C | S81 | E80 | K42 |
| PtGSTF1 | S69 | E68 | Q42, K43, Q55 |
| hLyso | R98 | Y63 | / |

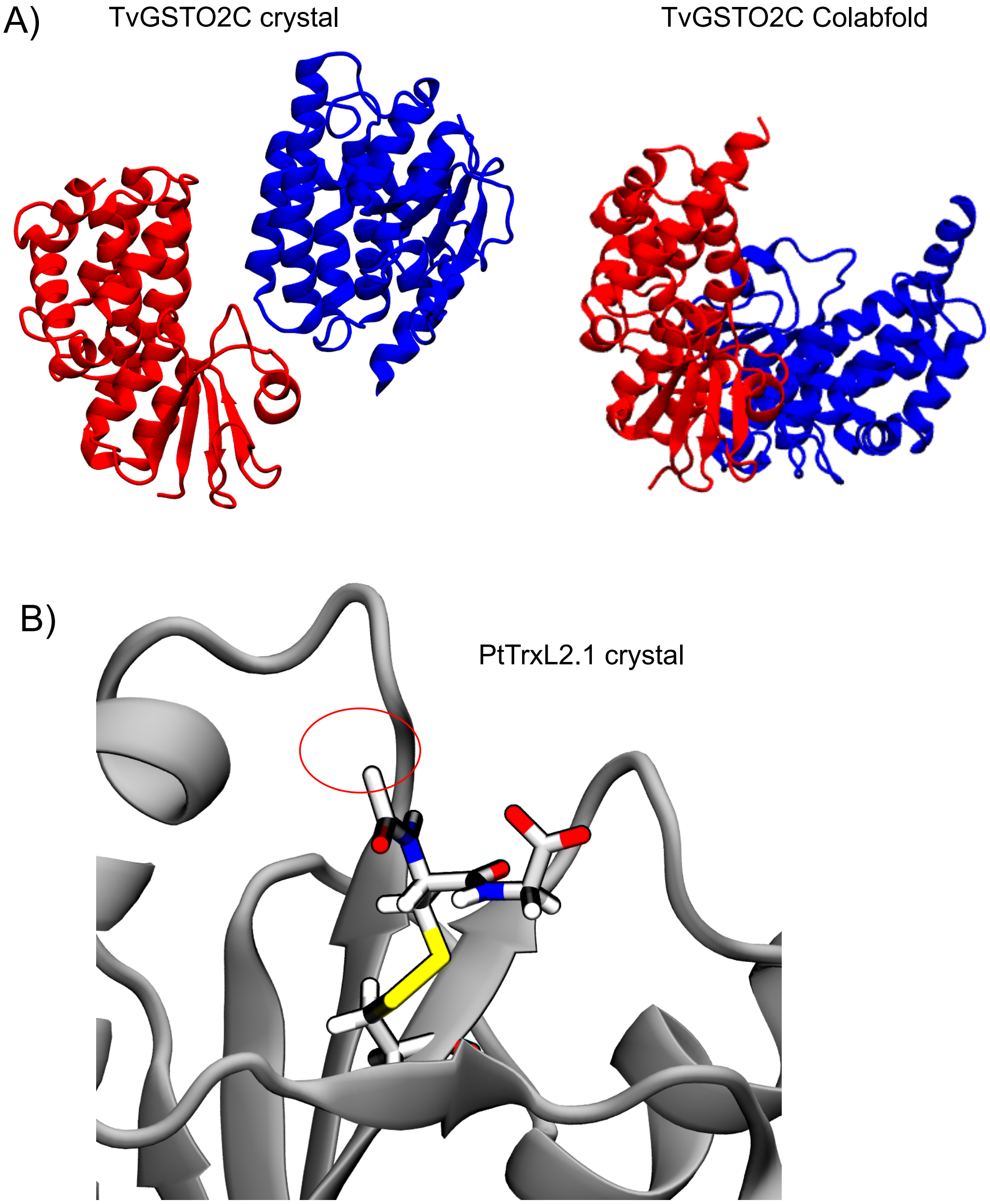


Figure S1 – A) Quaternary structure of the TvGSTO2C (PDB 6SR8) protein as displayed from the experimental PDB file (left) and the Colabfold prediction (right). B) CSG structure as present in the PtTrxL2.1 reference structure (PDB 5NYN), highlighting the absence of the terminal atoms on the glutamate side.

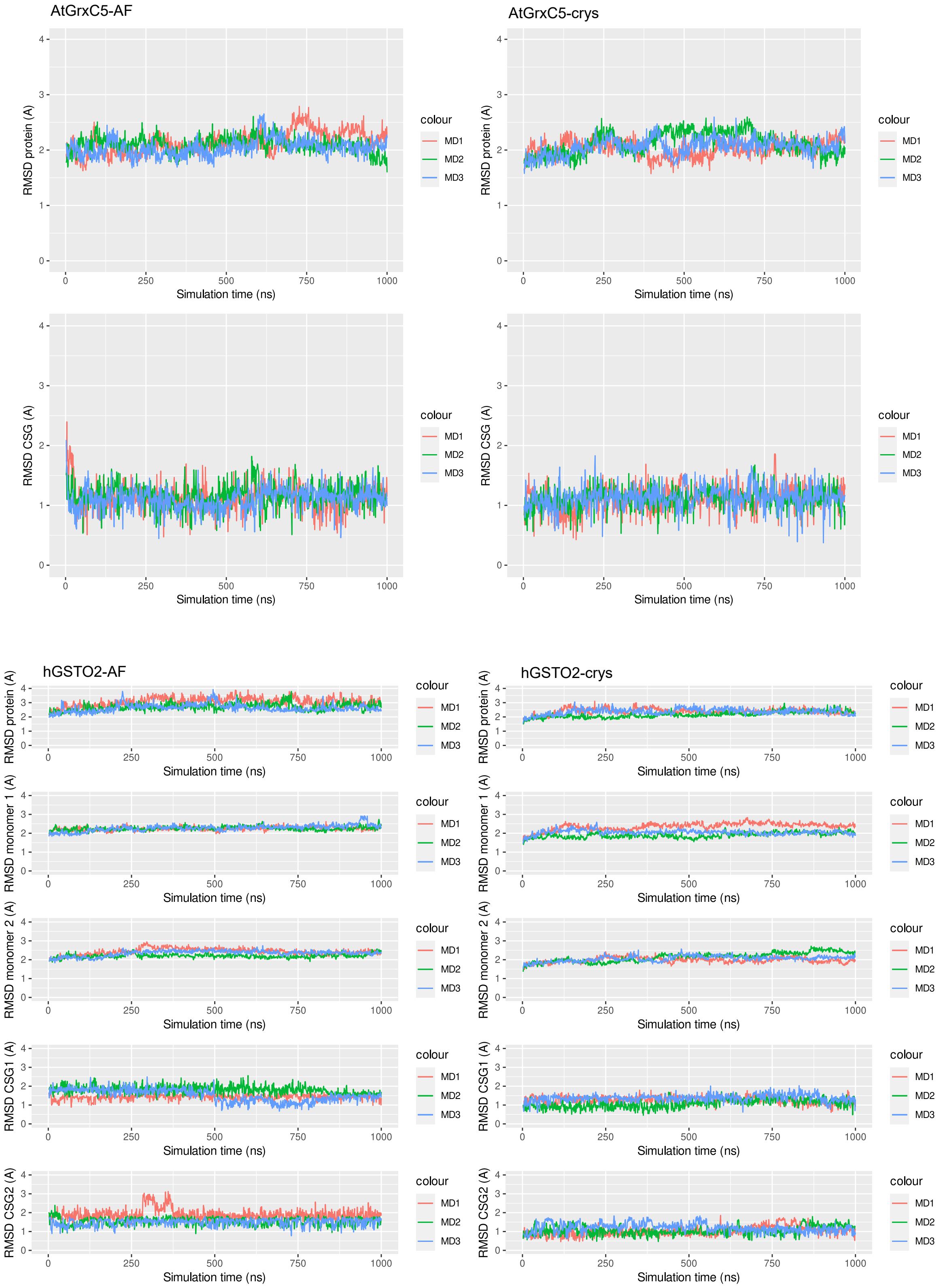


Figure S2 – Time series of the RMSD values (protein all atoms, protein backbone atoms, CSG residue) for the three replicates carried out in AF (left) and crys (right) simulations, for AtGrxC5 and hGSTO2.

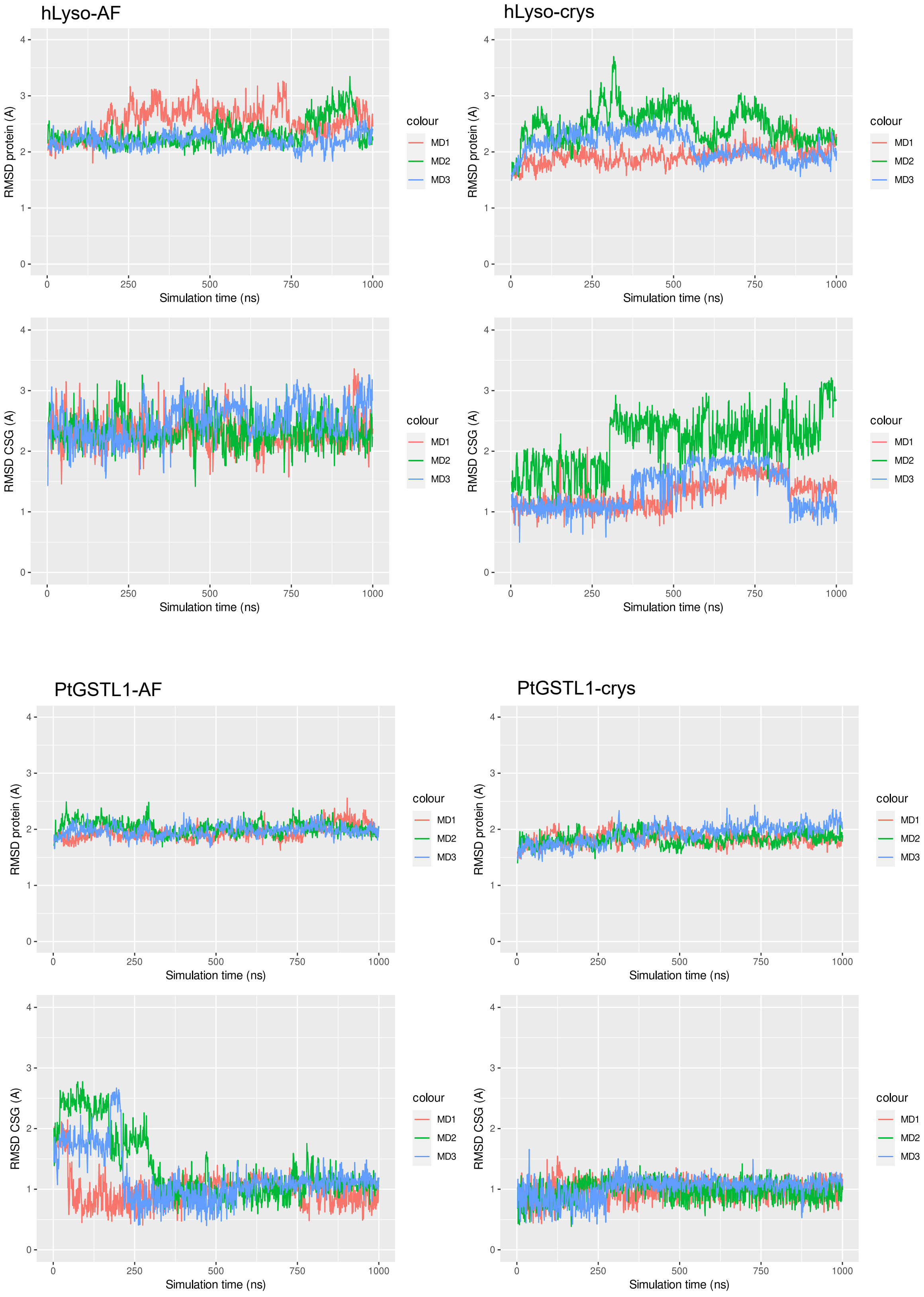


Figure S3 – Time series of the RMSD values (protein all atoms, protein backbone atoms, CSG residue) for the three replicates carried out in AF (left) and crys (right) simulations, for hLyso and PtGSTL1.

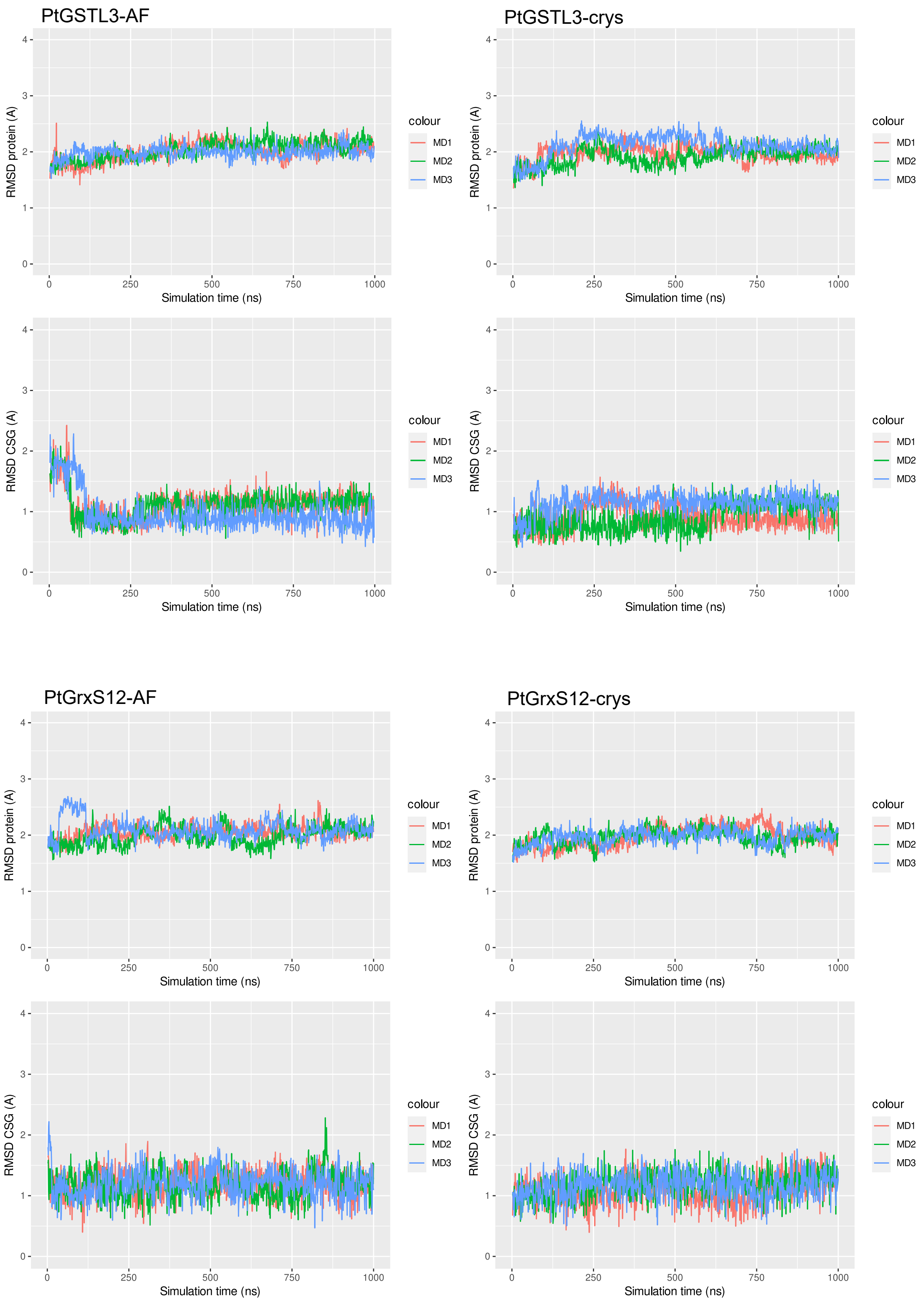


Figure S4 – Time series of the RMSD values (protein all atoms, protein backbone atoms, CSG residue) for the three replicates carried out in AF (left) and crys (right) simulations, for PtGSTL3 and PtGrxS12.

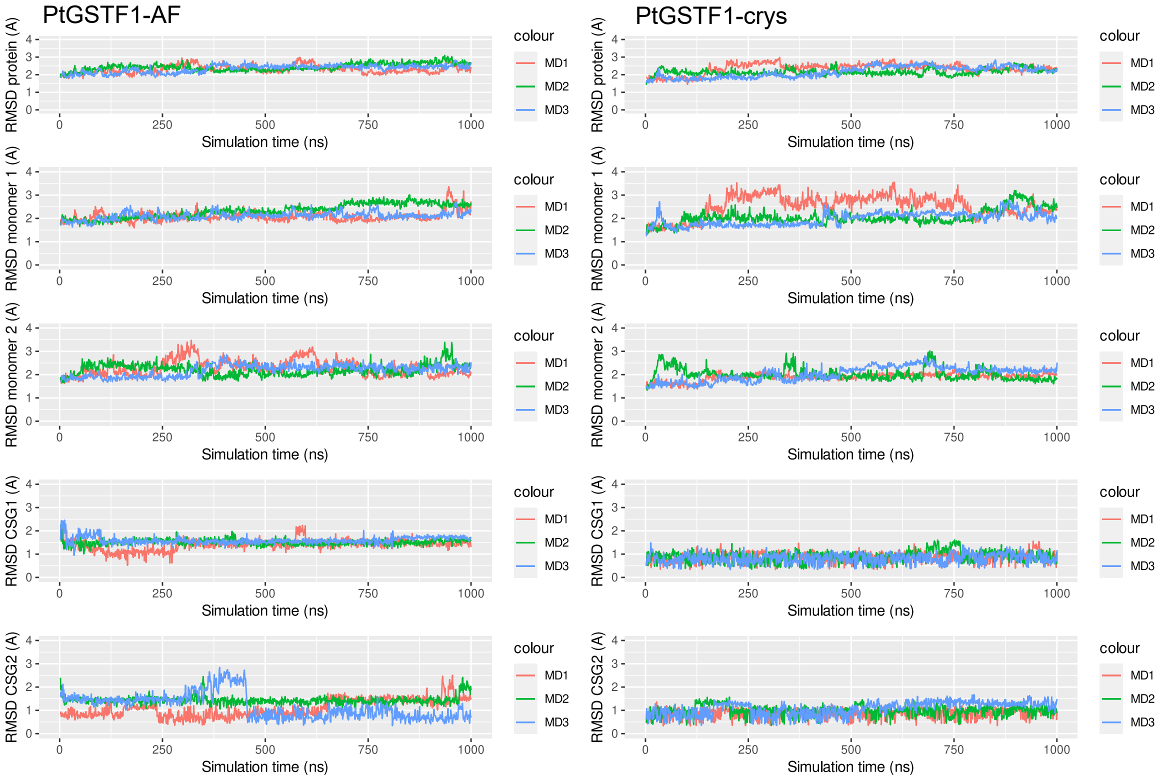


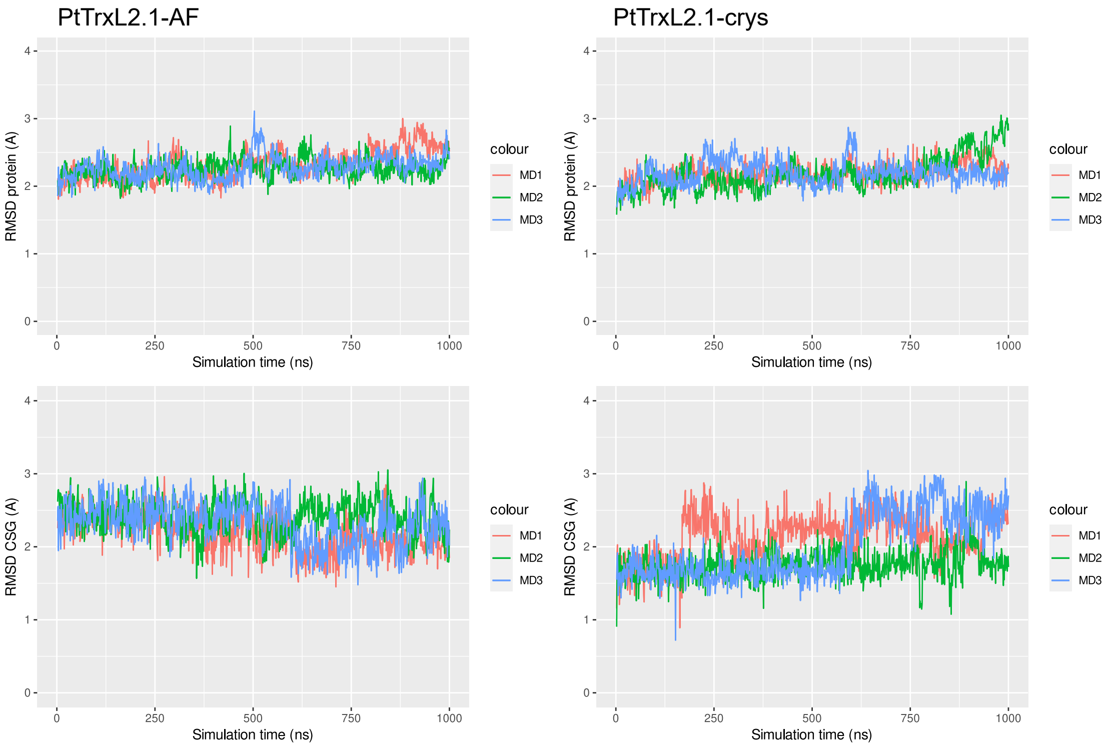


Figure S5 – Time series of the RMSD values (protein all atoms, protein backbone atoms, CSG residue) for the three replicates carried out in AF (left) and crys (right) simulations, for PtGSTF1 anf PtTrxL2.1.

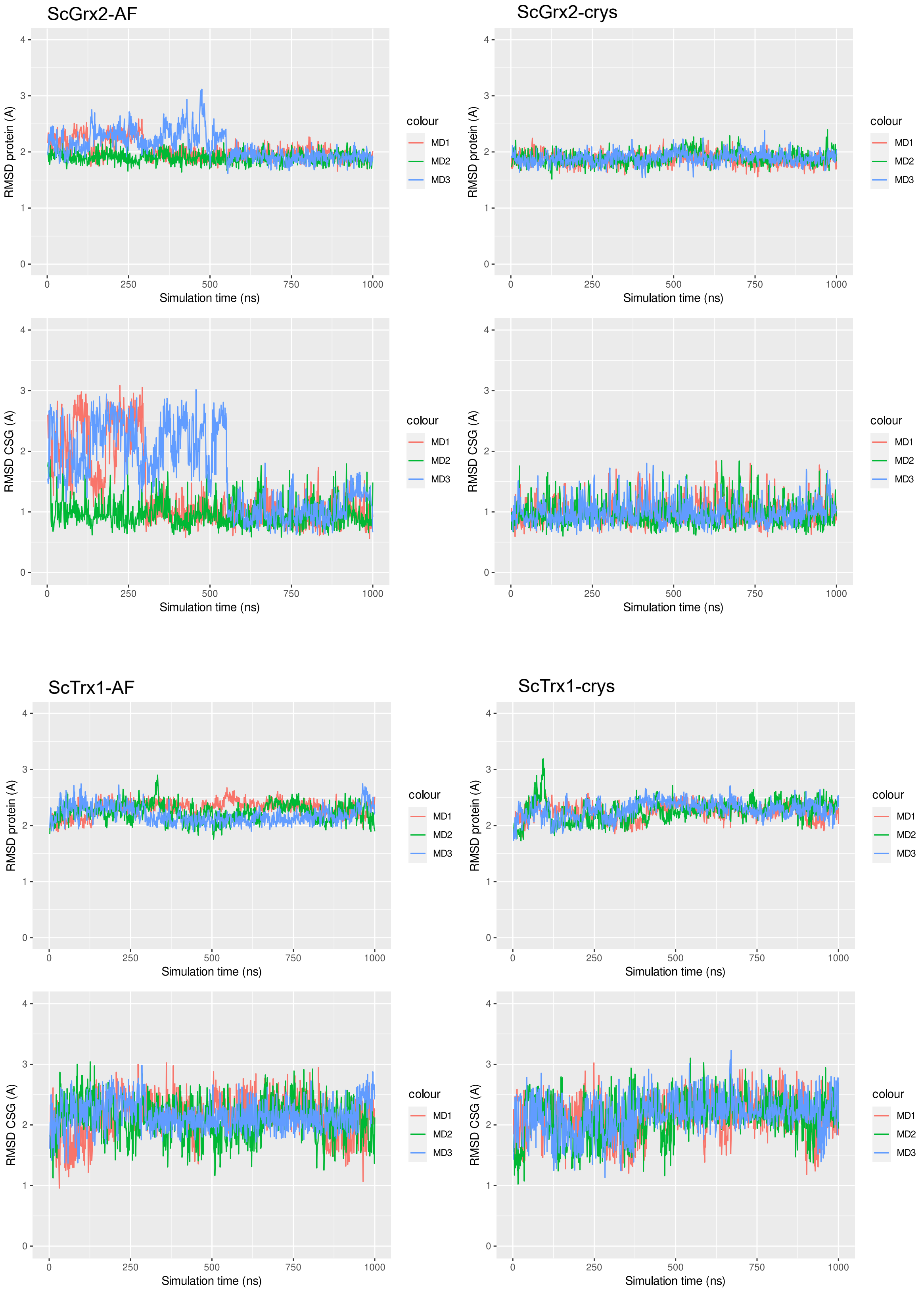


Figure S6 – Time series of the RMSD values (protein all atoms, protein backbone atoms, CSG residue) for the three replicates carried out in AF (left) and crys (right) simulations, for ScGrx2 and ScTrx1.

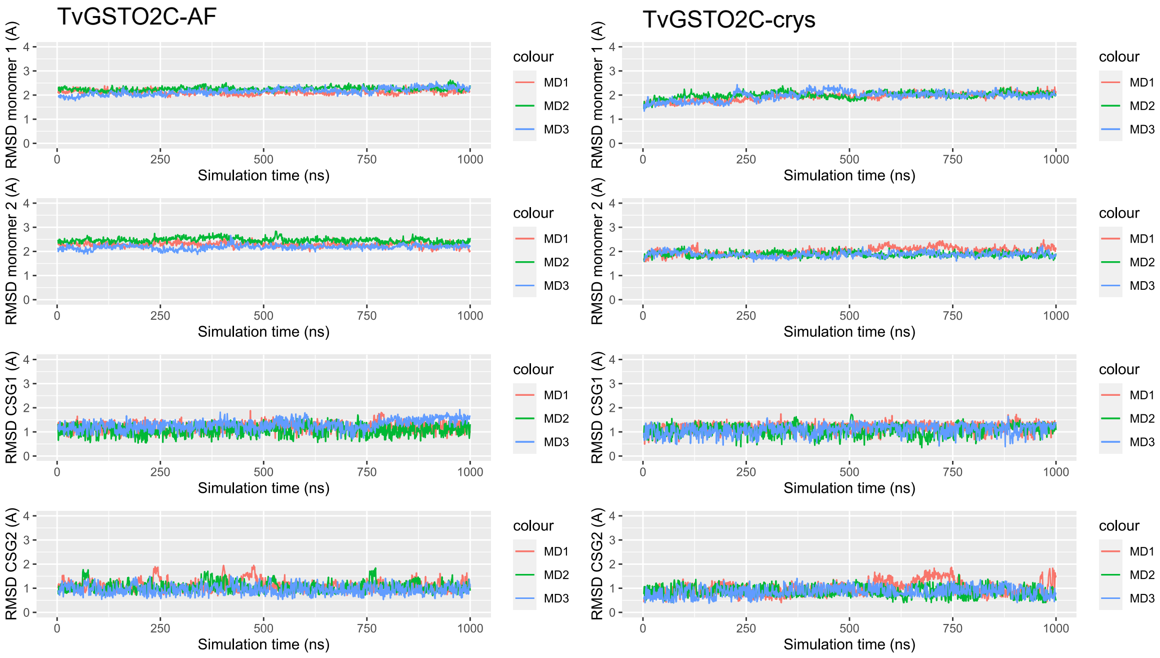


Figure S7 – Time series of the RMSD values (protein all atoms, protein backbone atoms, CSG residue) for the three replicates carried out in AF (left) and crys (right) simulations, for TvGSTO2C.

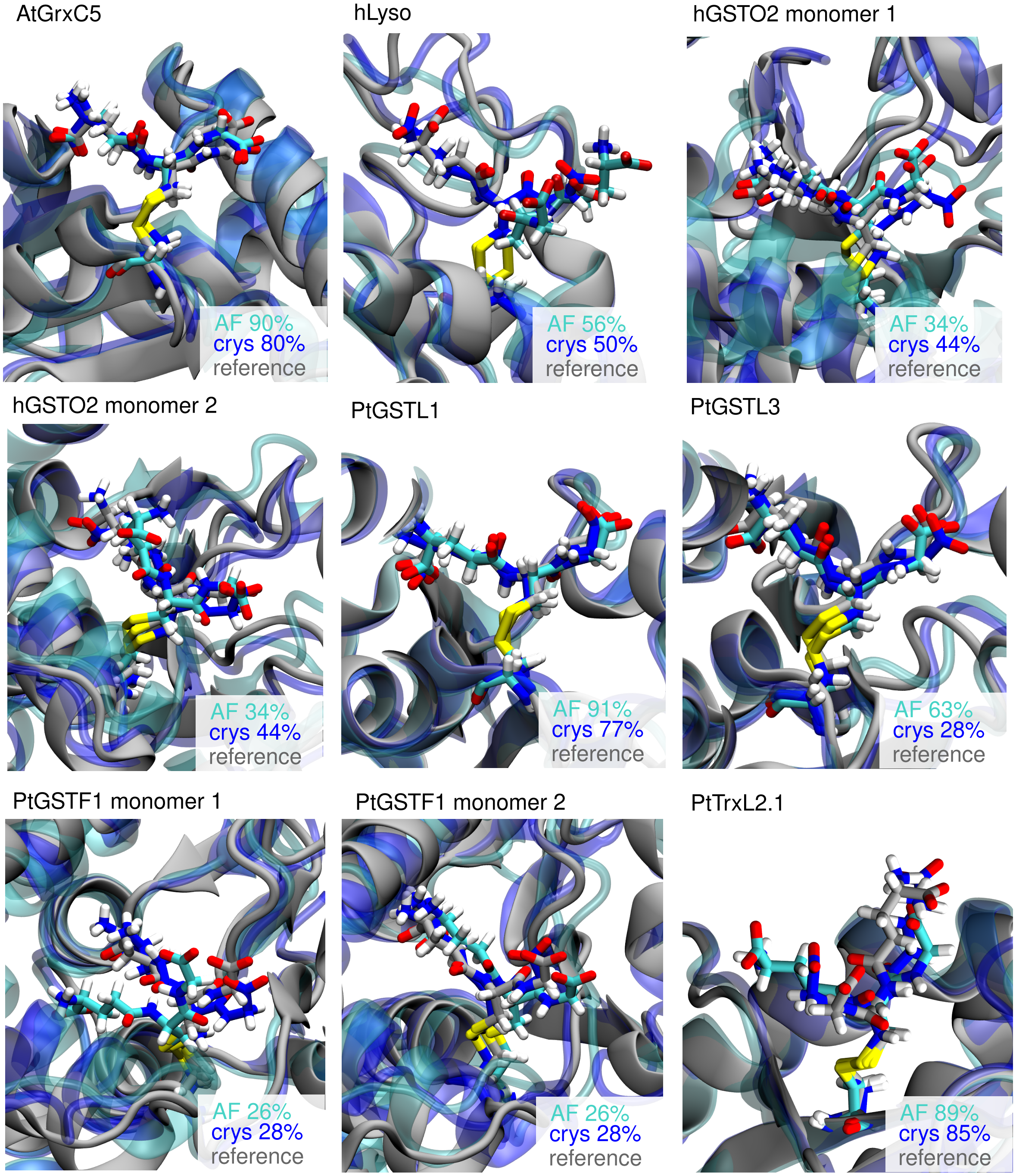


Figure S8 – Superimposition of representative structures of the major conformations of CSG as calculated from the cluster analysis of MD ensembles with the reference structure. Structures of the crystal reference and from the AF and crys simulations are displayed in gray, cyan and blue respectively. The percentage of occurrence of the clusters are also given.

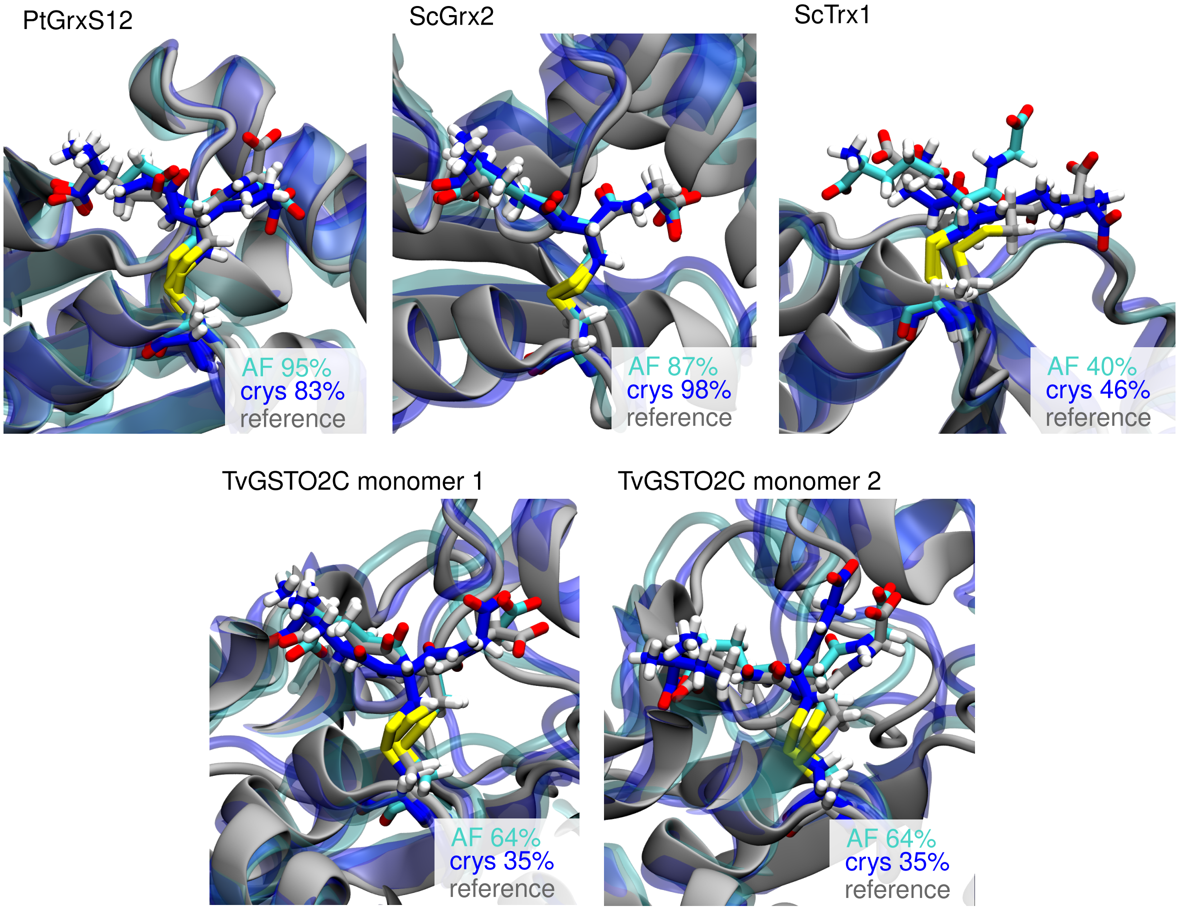


Figure S9 – Superimposition of representative structures of the major conformations of CSG as calculated from the cluster analysis of MD ensembles with the reference structure. Structures of the crystal reference and from the AF and crys simulations are displayed in gray, cyan and blue respectively. The percentage of occurrence of the clusters are also given.

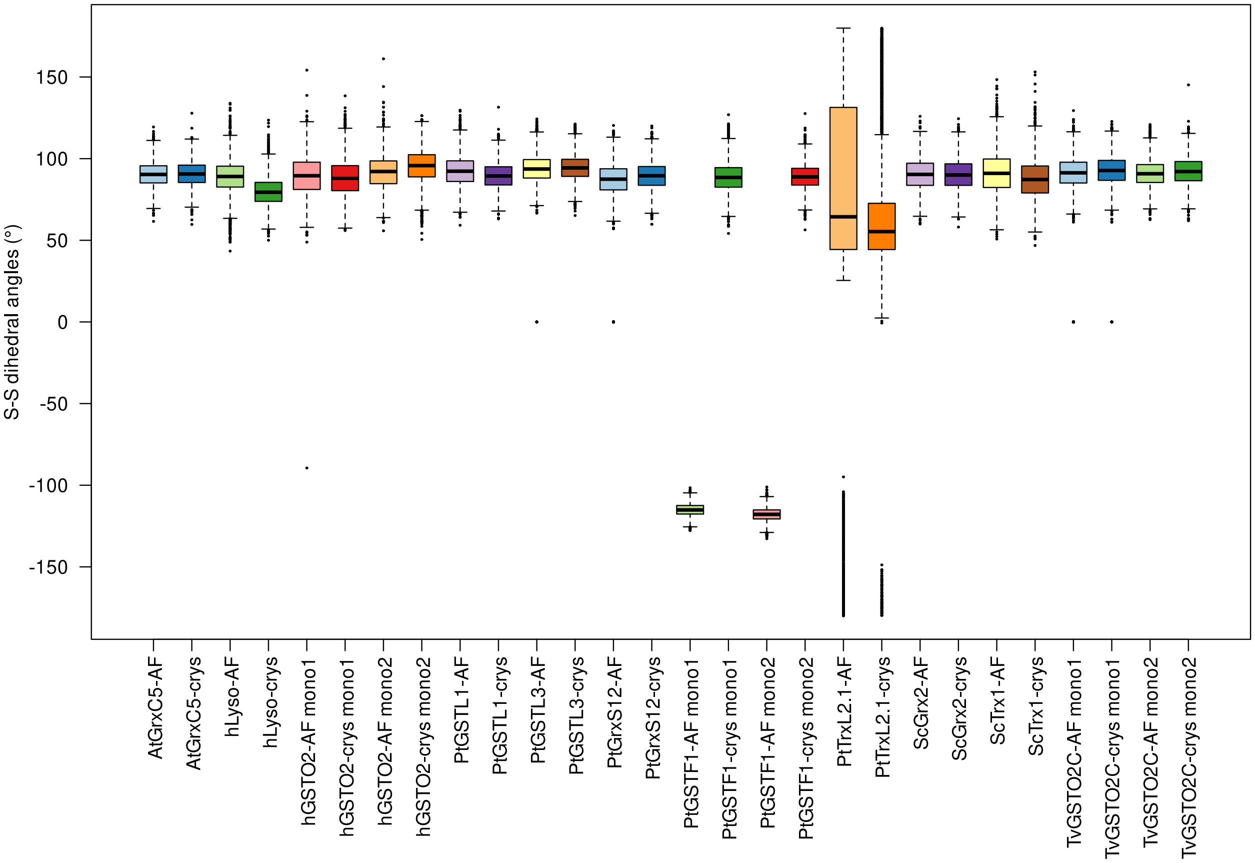


Figure S10 – Boxplot of CSG S-S dihedral angle values (°) in crys and AF simulation, for each system.

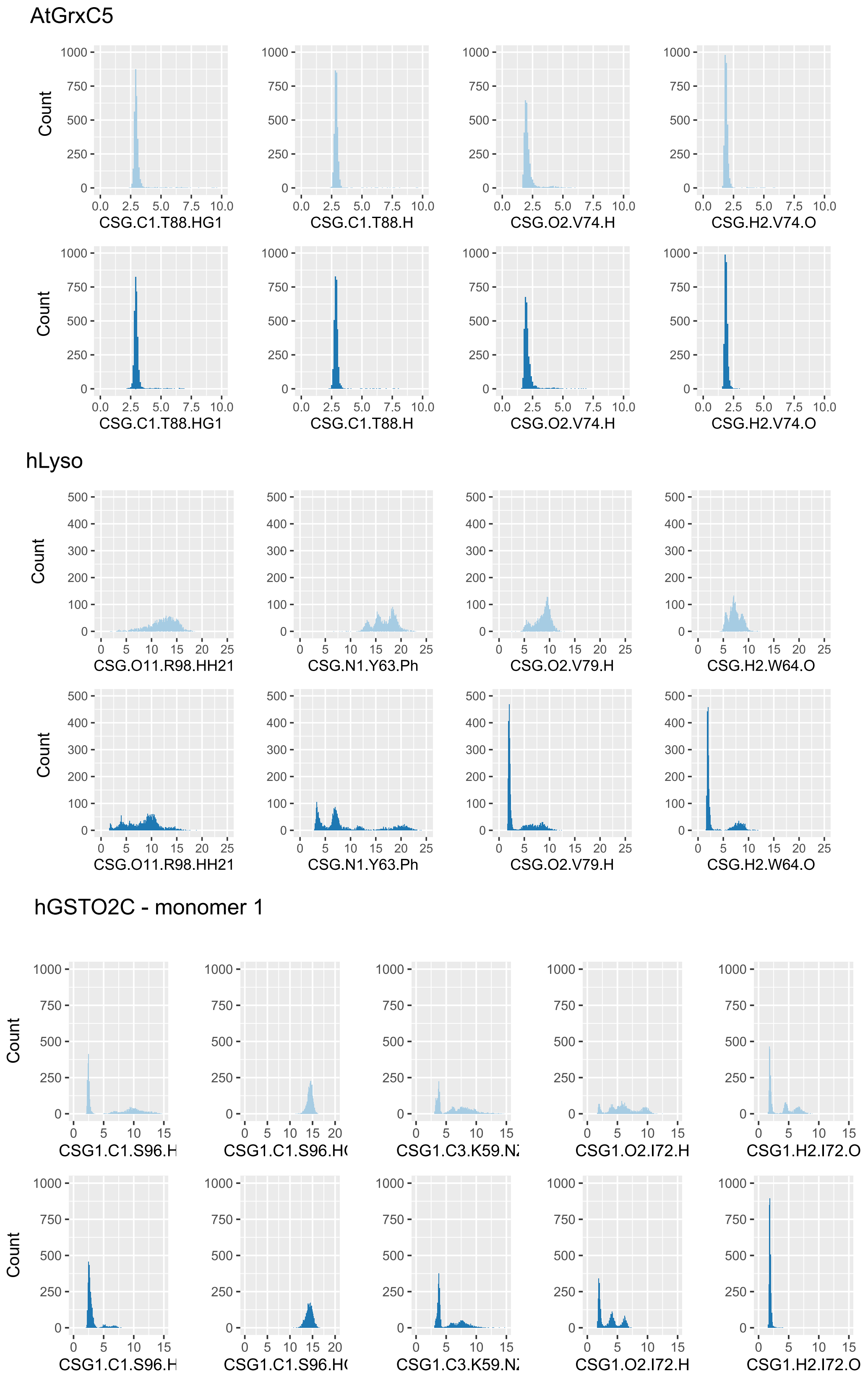


Figure S11 – Distribution of the key-distances involved in the native CSG interaction network, for AF (cyan) and crys (blue) simulations.

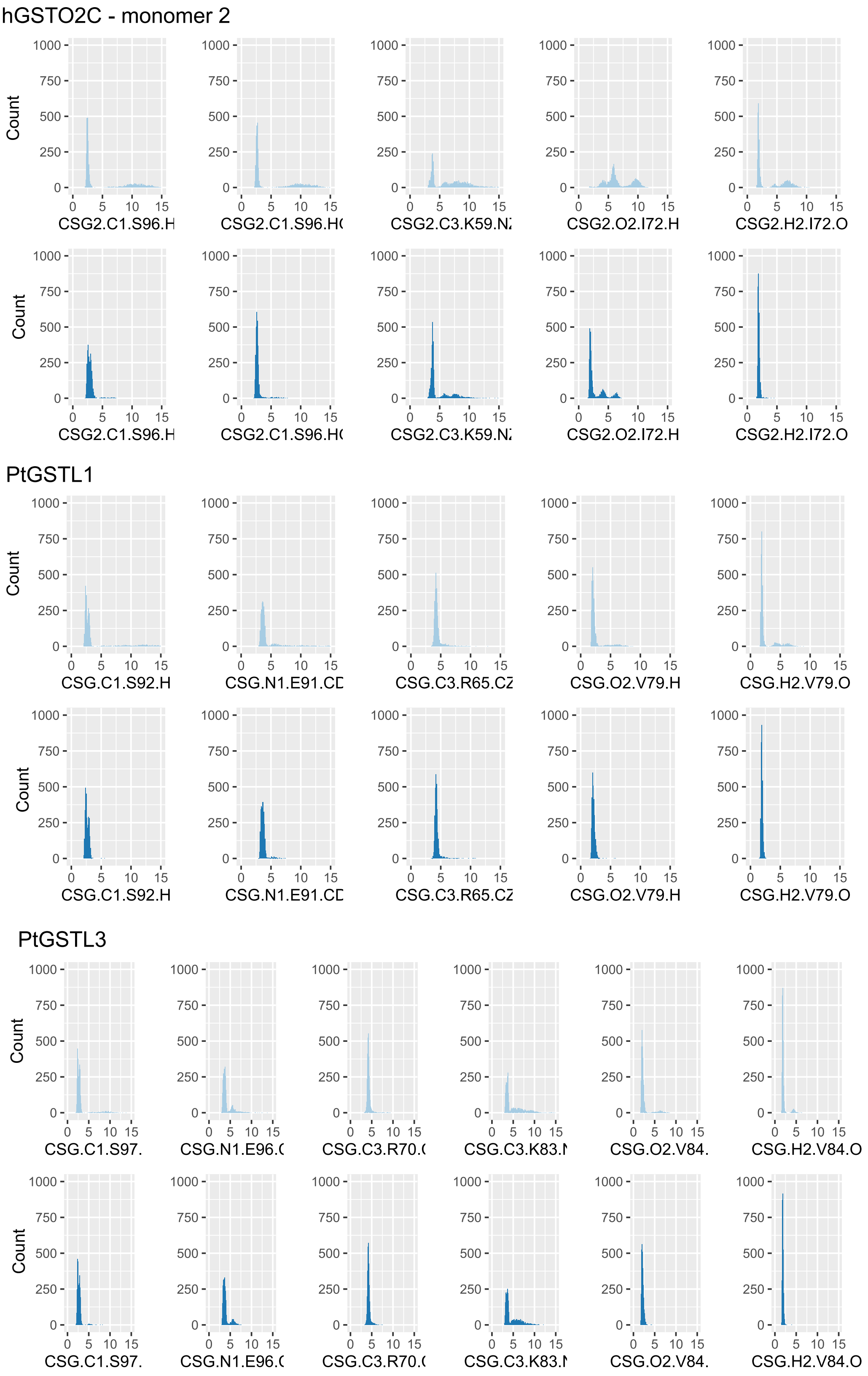


Figure S12 – Distribution of the key-distances involved in the native CSG interaction network, for AF (cyan) and crys (blue) simulations.

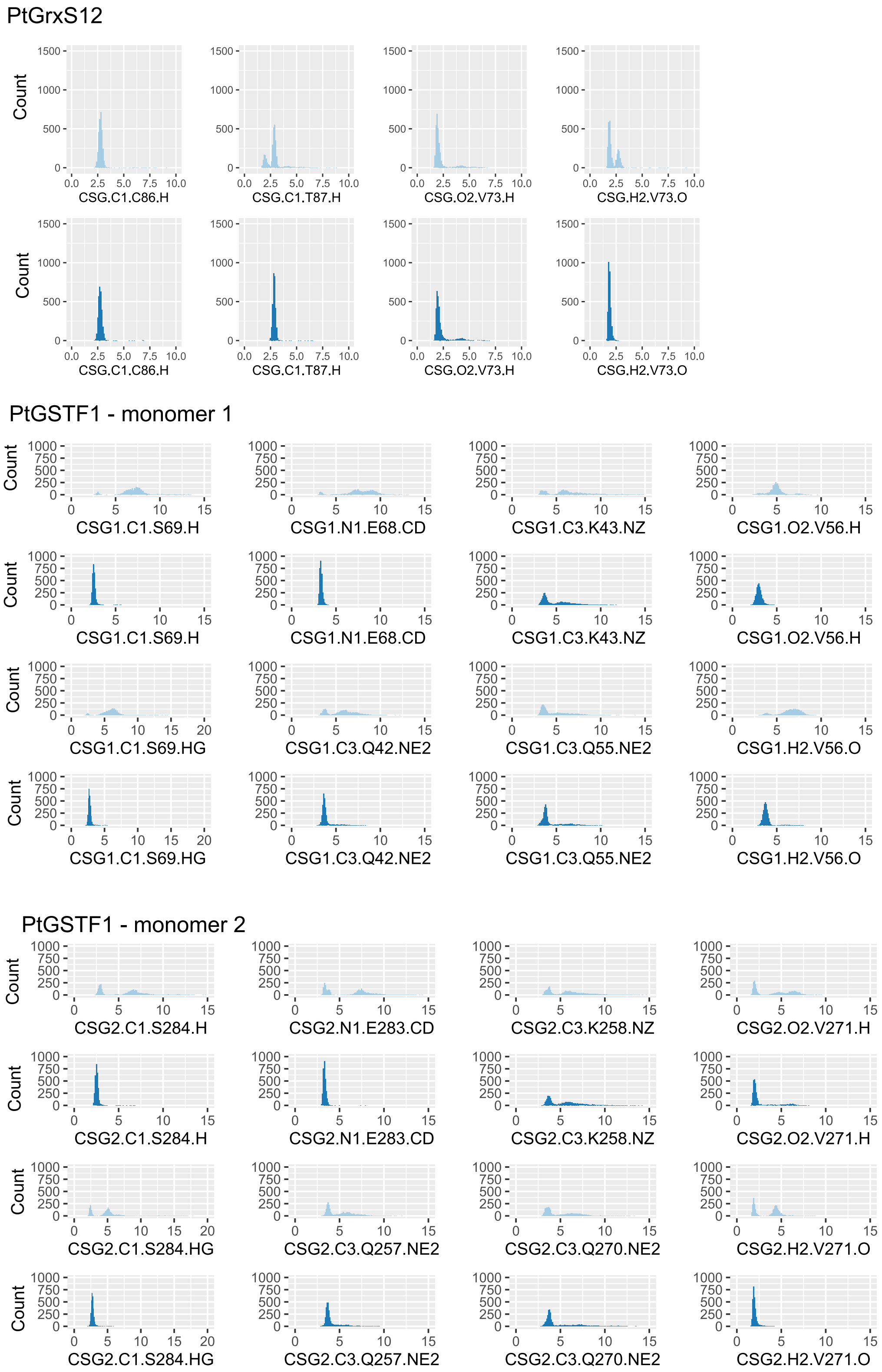


Figure S13 – Distribution of the key-distances involved in the native CSG interaction network, for AF (cyan) and crys (blue) simulations..

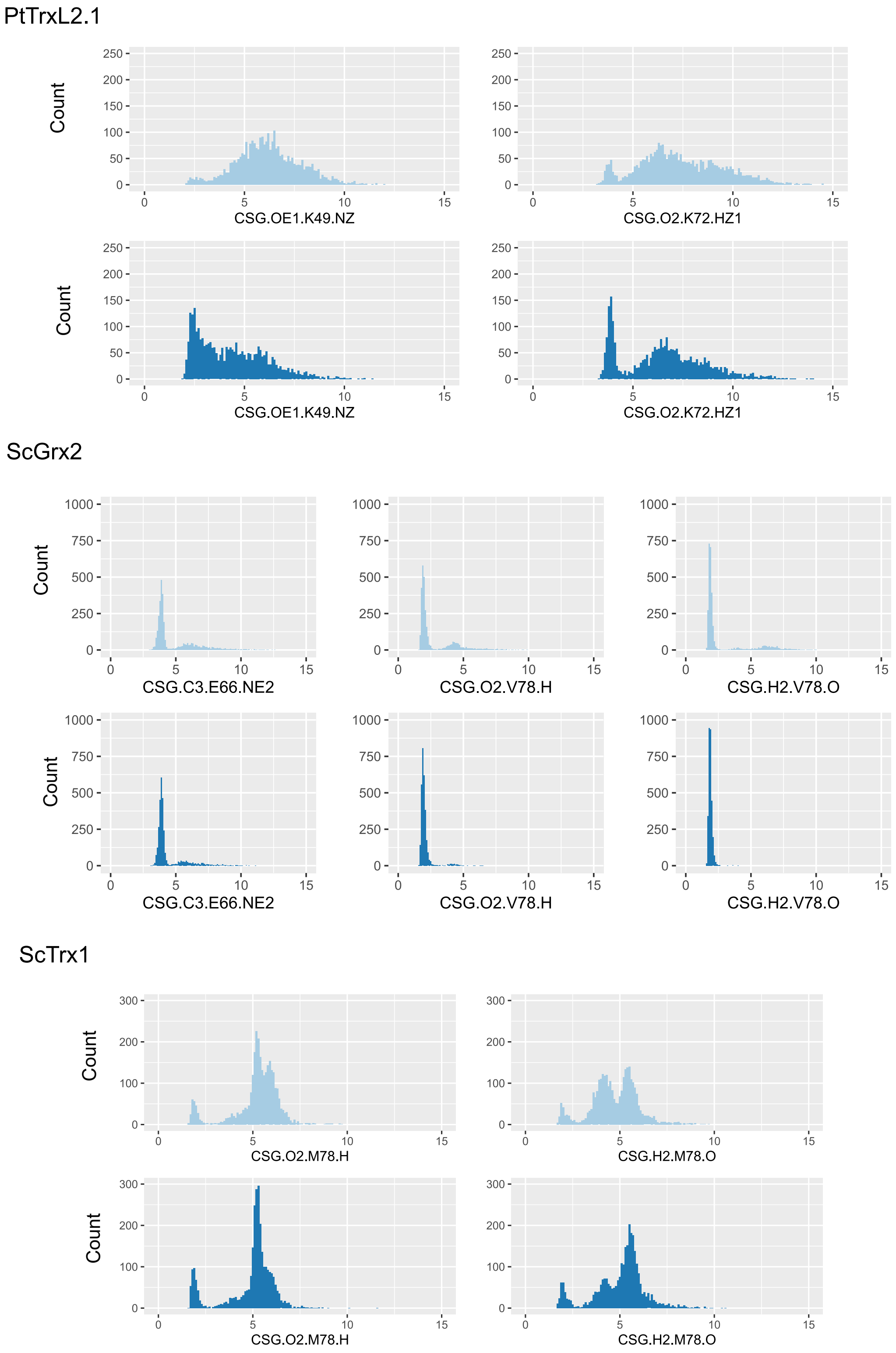


Figure S14 – Distribution of the key-distances involved in the native CSG interaction network, for AF (cyan) and crys (blue) simulations.

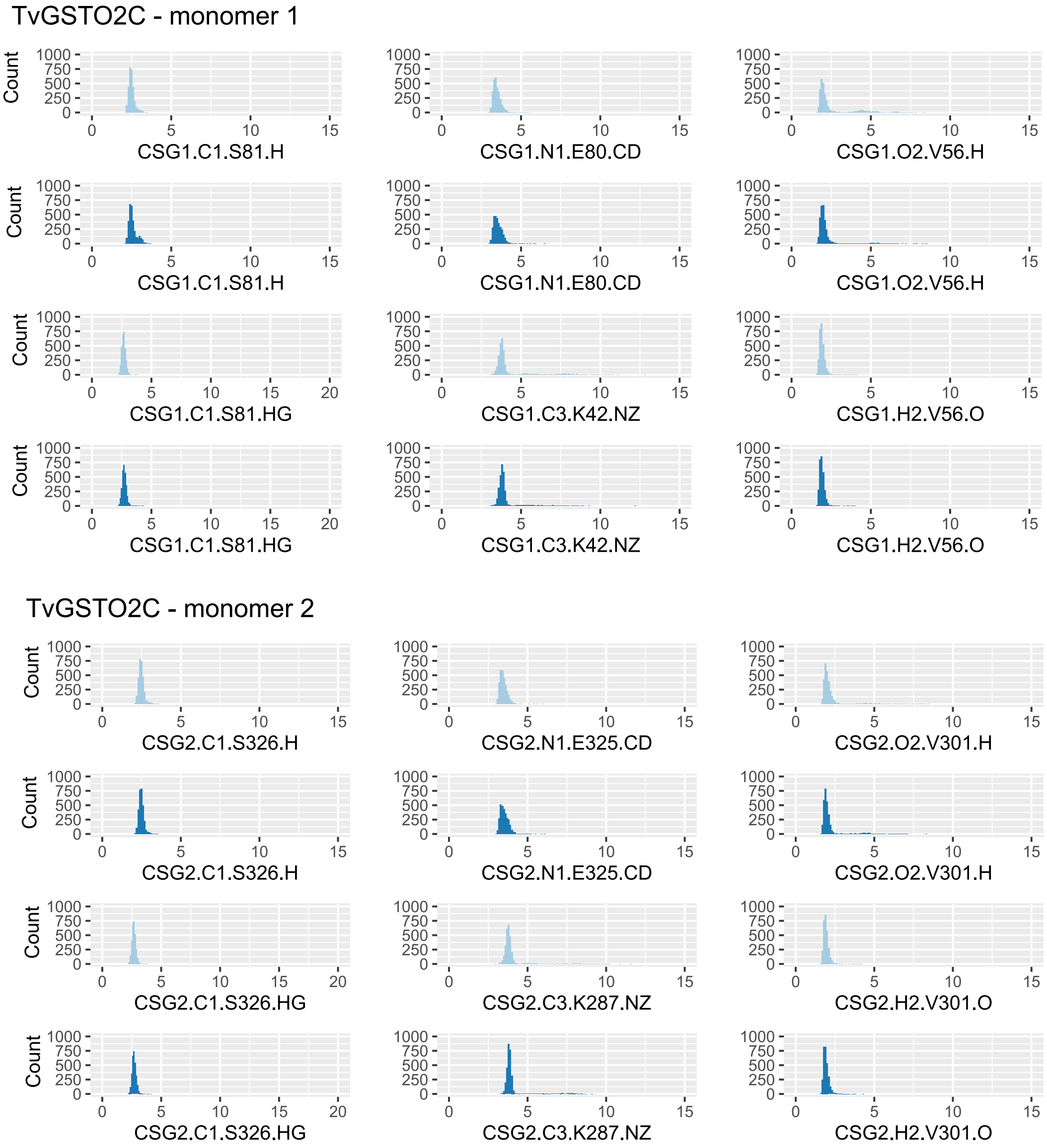


Figure S15 – Distribution of the key-distances involved in the native CSG interaction network, for AF (cyan) and crys (blue) simulations.

(6) Lallement, P.-A.; Meux, E.; Gualberto, J. M.; Prosper, P.; Didierjean, C.; Saul, F.; Haouz, A.; Rouhier, N.; Hecker, A. Structural and enzymatic insights into Lambda glutathione transferases from Populus trichocarpa, monomeric enzymes constituting

an early divergent class specific to terrestrial plants. *Biochemical Journal* **2014**, *462*, 39–52.

(7) Zhou, H.; Brock, J.; Liu, D.; Board, P. G.; Oakley, A. J. Structural insights into the dehydroascorbate reductase activity of human omega-class glutathione transferases. *Journal of molecular biology* **2012**, *420*, 190–203.

(8) Pégeot, H.; Koh, C. S.; Petre, B.; Mathiot, S.; Duplessis, S.; Hecker, A.; Didierjean, C.; Rouhier, N. The poplar Phi class glutathione transferase: expression, activity and structure of GSTF1. *Frontiers in Plant Science* **2014**, *5*, 712.

(9) Perrot, T.; Schwartz, M.; Deroy, A.; Girardet, J.-M.; Kohler, A.; Morel-Rouhier, M.; Favier, F.; Gelhaye, E.; Didierjean, C. Diversity of Omega Glutathione Transferases in mushroom-forming fungi revealed by phylogenetic, transcriptomic, biochemical and structural approaches. *Fungal Genetics and Biology* **2021**, *148*, 103506.

(10) Inaka, K.; Miki, K.; Kikuchi, M.; Taniyama, Y.; Matsushima, M. Structure of a glutathionylated human lysozyme: a folding intermediate mimic in the formation of a disulfide bond. *Acta Crystallographica Section D: Biological Crystallography* **1995**, *51*, 619–625.
